# Genomic variation in the tea leafhopper reveals the basis of adaptive evolution

**DOI:** 10.1101/2021.11.23.469662

**Authors:** Qian Zhao, Longqing Shi, Weiyi He, Jinyu Li, Shijun You, Shuai Chen, Jing Lin, Yibin Wang, Liwen Zhang, Guang Yang, Liette Vasseur, Minsheng You

**Author notes:** Correspondence should be addressed to Minsheng You. These authors contributed equally to this work: Qian Zhao and Longqing Shi.

## Abstract

The tea green leafhopper (TGL), *Empoasca onukii*, is of biological and economic interest. Despite numerous studies, the mechanisms underlying its adaptation and evolution remain enigmatic. Here, we used previously untapped genome and population genetics approaches to examine how this pest so rapidly has adapted to different environmental variables and thus has expanded geographically. We complete a chromosome-level assembly and annotation of the *E. onukii* genome, showing notable expansions of gene families associated with adaptation to chemoreception and detoxification. Genomic signals indicating balancing selection highlight metabolic pathways involved in adaptation to a wide range of tea varieties grown across ecologically diverse regions. Patterns of genetic variation among 54 *E. onukii* samples unveil the population structure and evolutionary history across different tea-growing regions in China. Our results demonstrate that the genomic change in key pathways, including those linked to metabolism, circadian rhythms and immune system function, may underlie the successful spread and adaptation of *E. onukii*. This work highlights the genetic and molecular bases underlying the evolutionary success of a species with broad economic impact, and provides insight into insect adaptation to host plants, which will ultimately facilitate more sustainable pest management.

## Introduction

Tea is the most popular beverage worldwide, surpassing coffee and cocoa, with a production of 6.1 million metric tons in 2019 (ITC; https://www.statista.com/statistics/264183/global-production-and-exports-of-tea-since-2004/). China represents the largest tea producer, consumer, and exporter in the world. In Asia, the tea green leafhopper (TGL), *Empoasca onukii* (Hemiptera: Cicadellidae), represents the most devastating pest across tea plantations, causing up to 50% economic loss of tea production annually [1, 2]. Both nymph and adult TGLs pierce and suck the sap of tender tea shoots, which are the most important part of the plant to produce high-quality tea. Adult females also lay their eggs in these shoots, leading to irreparable damage (Figure S1) [2, 3]. Presence of local TGL populations has been recorded in China since the 1950s [4]. Its distribution has increased around tea-producing regions of China, Japan, and Vietnam [5]. *E. onukii* can cause yield loss of between 15 and 50%, up to 100% in severely damaged plantations [2, 6].

*E onukii* belongs to the most species-rich hemimetabolous order, various species of which are agricultural pests or human disease vectors [7]. As a monophagous insect, TGL is well-adapted, both physiologically and biochemically, to different tea varieties [8]. Thus, the rapid expansion of *E. onukii* raises critical questions concerning which factors contribute to its successful dispersal and colonization, and how genomic architecture underlies its broad and rapid ability to adapt.

To address the above questions, we generated a chromosomal level genome assembly of the *E. onukii* by integrating Illumina short reads, Oxford Nanopore long reads, and high-throughput chromosome conformation capture (Hi-C technology). This high-quality genome resource enabled us to investigate the genetic basis of chemoreception and detoxification in this insect, key to adapting to new environments. Based on 54 re-sequenced genomes of the *E. onukii* samples collected from different locations across a diverse range of tea-growing regions in China, we analyzed patterns of genomic variation and population structure in this species, allowing us to gain insights into its evolutionary history and successful, rapid spread and colonization.

## Results and Discussion

### Chromosome-level assembly of the tea green leafhopper

The genome of *E. onukii* was estimated to be ∼608Mb based on K-mer analysis. We combined 61× Illumina short read and 109× Nanopore ONT sequences with chromosome-scale scaffolding. We informed our assembly using physical mapping of high-throughput chromatin conformation capture (Hi-C) (Tables S1 and S2), to generate an assembly based on 599 Mb of sequence, with the mitochondrial sequences excluded (**Table 1** and Tables S3, S4). This assembly accounted for 98.5% of the estimated genome size. A total of 592 Mb sequences and 98.83% of assembled sequences were then anchored onto 10 pseudo-chromosomes using ALLHiC (see Methods and **Tables 1 and 2**, Figures 1A and S2A). An official gene set was generated based on alignment of insect gene homologs, *ab initio* predictions, and transcriptomic evidence. Genome annotation predicted 19,642 protein coding genes with 92.5% BUSCO completeness in TGLs (Table 1 and S5). The sequenced *E. onukii* genome showed high heterozygosity (2.8%), with 13,122,207 heterozygous SNPs, 3,796,369 heterozygous indels, and complex segmental duplication patterns (**Figure 1**A). We also assembled the mitochondrial genome, which had a total length of 14.2 kb and 13 protein-coding genes annotated (Figure S2B).

**Figure 1.**
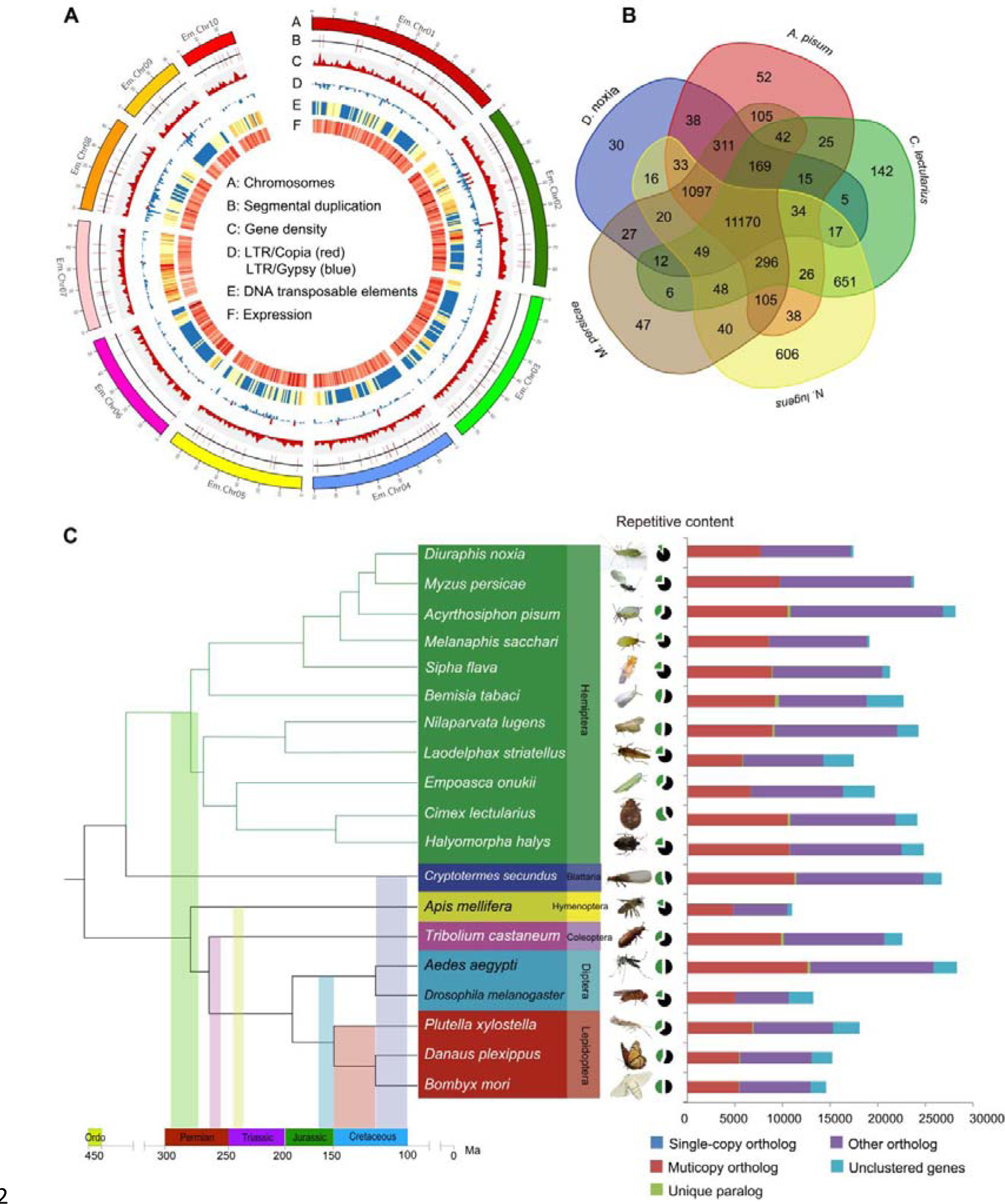
Genomic characterization of *E. onukii* and comparison with other insect genomes. **(A)** Genomic characterization of the sequenced *E. onukii*. The circles (from outermost to innermost) represent monoploid genome in Mb, segmental duplication, gene density, LTR Copia/Gypsy, DNA transposable elements and expression profiles. (B) Numerical comparison of homologous genes between gene sets from *E. onukii* and each of the five Hemiptera species. Dataset overlaps were determined using a BLASTP search (*e* value < 10^-5^). (C) Phylogenetic relationships among 15 insect species based on genomic comparisons. Single copy orthologs: only one copy in different genomes, multicopy orthologs: more than one copy in different genomes, unique paralogs: species-specific genes, other orthologs: unclassified orthologs, unclustered genes: genes that cannot be clustered into known gene families. Details about the identification are previously described [73].

**Table 1.** Sequencing, chromosome-scale assembly and annotation of the *E. onukii* genome.

| **Sequencing** |  |
| --- | --- |
| Sequencing Platform | Nanopore ONT |
| Data size (Gb) | 65 |
| Genome sequencing depth (×) | 109 |
| Estimated genome size (Mb) | ~608 |
| **Chromosome-scale assembly** |  |
| Assembly size (Mbp) | 599 |
| % of estimated genome size | 98.5 |
| No. of contigs | 1800 |
| Contig N50 (Mb) | 2.2 |
| Average length (bp) | 332,835 |
| Minimum contig length (bp) | 2,552 |
| No. of chromosomes | 10 |
| Scaffold N50 (Mb) | 67.98 |
| No. of unanchored contigs | 234 |
| Length of anchored contigs (Mb) | 592 |
| Anchor rate (%) | 98.83 |
| BUSCO completeness (%) | 92.7 |
| **Annotation** |  |
| No. of protein-coding genes | 19,642 |
| Average gene length (bp) | 7904 |
| Average CDS length (bp) | 201 |
| Average exon number per gene | 4.99 |
| BUSCO completeness (%) | 92.5 |

**Table 2.** Chromosome-based statistics of the *E. onukii* genome.

| Chromosome | No. of contigs | Length (bp) |
| --- | --- | --- |
| Chr1 | 263 | 94,216,414 |
| Chr2 | 314 | 91,501,843 |
| Chr3 | 121 | 74,646,503 |
| Chr5 | 202 | 67,983,564 |
| Chr4 | 155 | 65,611,511 |
| Chr6 | 79 | 54,167,312 |
| Chr8 | 177 | 48,146,360 |
| Chr7 | 88 | 43,255,712 |
| Chr9 | 110 | 28,073,217 |
| Chr10 | 57 | 24,627,475 |
| Total number of contigs | 1800 |  |
| Total length of contigs (bp) | 599,103,029 |  |
| Total number of anchored contigs | 1566 |  |
| Total length of chromosome level assembly (bp) | 592,229,911 |  |
| Number of unanchored contigs | 234 |  |
| Length of unanchored contigs | 7,028,718 |  |
| Anchor rate (%) | 98.83 |  |

**Table 3.**
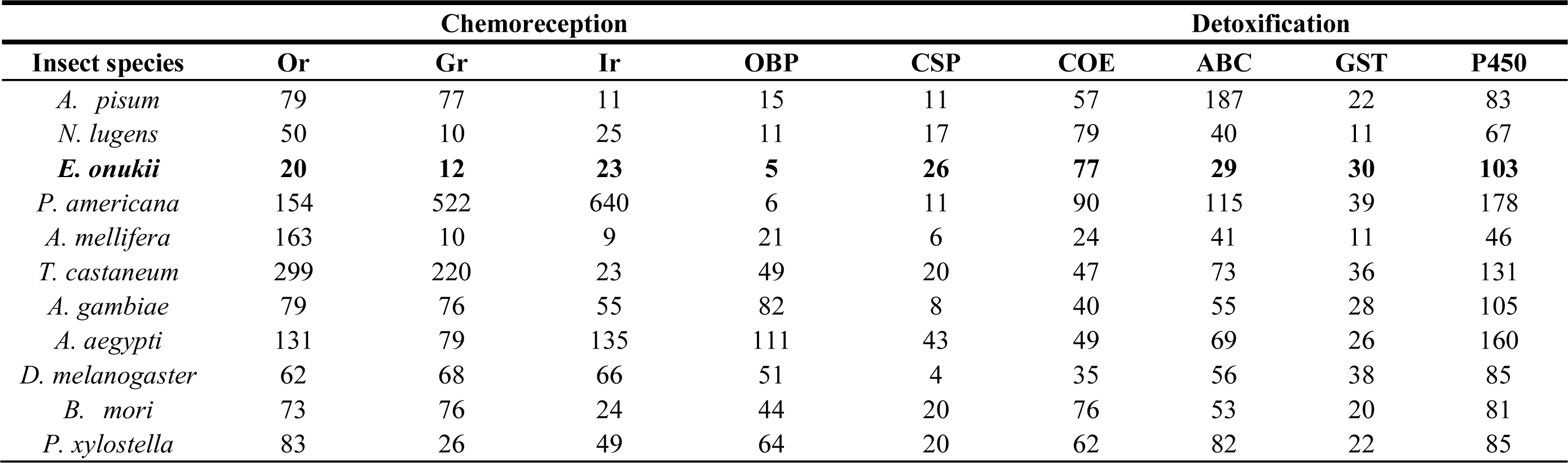
Numerical comparison of genes related to chemoreception and detoxification among different insect species.

Compared to recently published Hemiptera genomes [9, 10], this assembly is a high-quality genome with 92.7% BUSCO completeness (Tables S4 and S6). Around 92.3% (337.5/365.8 million) of the Illumina short reads were mapped to the assembled reference, representing about ∼94% of the genome (Table S7). Well-organized patterns of interacting contact along the diagonal for each pseudo-chromosome confirmed the high-quality chromosome-level assembly (Figure S2). In addition, assessment using the LTR Assembly Index (LAI) [11] revealed that more intact LTRs were recalled in our assembly than previously published insect genomes [9, 10], further supporting the high quality of the TGL genome (Figure S3A).

**Table 4.** Number of populations, nuclear SNPs, and genetic diversity (π) of the three clustered groups.

| **Group** | **No. populations** | **No. SNPs** | **No. Indels** | **$\pi$** |
| --- | --- | --- | --- | --- |
| Group I | 4 | 52,089,001 | 14,913,280 | 0.004062 |
| Group II | 28 | 472,824,888 | 135,096,045 | 0.004744 |
| Group III | 22 | 279,181,783 | 79,642,551 | 0.004662 |

In total, 19,642 genes were annotated in *E. onukii* and compared with five other well-annotated Hemiptera published genomes including *Diuraphis noxia* [12]*, Acyrthosiphon pisum* [13]*, Cimex lectularius* [9]*, Nilaparvata lugens* [14], and *Myzus persicae* (https://bipaa.genouest.org/sp/acyrthosiphon_pisum/). Results showed that 56.9% (11,170/19,642) of *E. onukii* genes had homologs (Figure 1B). The *E. onukii* genome contained ∼37.7% repetitive sequences, a relatively moderate level among published Hemiptera genomes, which repetitive sequences range from ∼12% in *D. noxia* [12] to 56.5% in *C. lectularius* [9]. TEs accounted for approximately 32.8% of the *E. onukii* genome, and were comprised of 12.82% transposon sequences, 4% long terminal repeats (LTRs), 11.24% long interspersed nuclear elements (LINEs), 2.20% short interspersed nuclear elements (SINEs) (Table S8). TE level in *E. onukii* was comparable to that of *N. lugens* [15], but approximately 1.5 times higher than *A. glycines* [10] (Figure 1C). Similar to the aphid genome, LINEs were more prevalent than LTR retroelements (Table S8). The moderate genome size and levels of repetitive sequences in *E. onukii* compared to other hemipteran species (Figure 1C; Figure S3B) indicated that TEs and other non-coding DNA might contribute to variation in genome size [16]. The genome sizes of insect generally result from changes in the repetitive DNA content [16]. Therefore, we calculated the correlation between repetitive sequence content and genome size in sequenced insect species from both the Holometabola (e.g., flies, beetles, wasps, and butterflies) and Hemimetabola (e.g., aphids, true bugs, blood-feeding bugs, and leafhoppers). As expected, the genome size was highly correlated with the DNA repetitive content (Spearman test, *r* = 0.8, *P* = 0.0004) (Figure S3B). Previous studies suggest that differences in DNA repeat content are likely due to TE variation or the influence of stochastic population effects [17].

We used OrthoFinder to identify orthologous genes across the genomes of *E. onukii* and other 18 insect species covering six different insect orders (Hemiptera, Isoptera, Hymenoptera, Coleoptera, Diptera and Lepidoptera). A total of 196 single-copy orthologous genes, 6411 multi-copied orthologous genes, 18 unique paralogous genes and 3325 unclustered genes were identified. The phylogenetic relationships among 19 sequenced insect species were analyzed using the PROTGAMMALGX model in RAxML [18] based on the 196 single-copy orthologous genes (Figure 1C). Based on these analyses, *E. onukii* was estimated to have diverged from *N. lugens* and *L. striatellus* approximately 175 Mya ago (Figure S4).

Expansion and contraction of gene families were analyzed based on 19 species. Results showed that both total and species-specific genes in Hemiptera genomes increased relative to other insect orders (Figure S4) [15]. We identified 2,859 novel genes (species-specific) in *E. onukii*, representing about 14.5% of the genome. In addition, 1178 expanded gene families were detected and these gene families were over-represented in specific Gene Ontology (GO) terms, including carboxylic ester hydrolase activity, zinc ion binding, iron ion binding, and transmembrane transporter activity (Table S9). The *E. onukii* genome contained 3880 contracted genes families (Figure S4). For example, functional analysis revealed that these genes were involved in immunity (immunoglobulins), myosin, and tropomyosin (Table S10). These *E. onukii*-specific gene family expansions and contractions were likely involved in evolutionary adaption to tea phloem sap, symbiotic dependence, pathogen immunity, and environmental conditions such as ecological and climatic variation, and tea variety differences. For example, evidence shows carboxylic ester hydrolase activity is involved in sap-sucking insects (e.g., *M. persicae* and *S. graminum*) sequestering OP insecticides [19]. Immunoglobulin superfamily proteins have been reported as candidates for synapse targeting functions related to synaptic specificity in the visual system in *Drosophila* [20, 21].

### Genomic adaptation to chemoreception and detoxification

The chemosensory system is essential for herbivorous insects to orient toward and locate potential host plants [22], potentially indicating how herbivorous insects adapt to host changes. Environmental signals and chemosensory stimuli are recognized and transduced by several multi-gene families including olfactory receptors (ORs), ionotropic receptors (IRs), gustatory receptors (GRs), odorant-binding proteins (OBPs), and chemosensory proteins (CSPs) [22, 23]. To examine genes linked to chemosensory stimuli recognition, we manually annotated several related gene families, including 20 olfactory receptors (ORs), 23 ionotropic receptors (IRs), 12 gustatory receptors (GRs), 5 odorant-binding proteins (OBPs), and 26 chemosensory proteins (CSPs) **(Table 3)**.

Comparative analysis of genomes across different species revealed an increased number of CSPs in the *E. onukii* genome (Table 3, **Figure 2**A and B, and Table S11). The phylogenetic analysis of Hemiptera identified 10 homologous subgroups of CSPs (CSP1-CSP10) (Figure 2B), which was consistent with a previous study [24]. Other than CSP5 and CSP6, *E. onukii* CSPs were present in seven of the ten clades, indicating these genes are highly conserved across the Hemiptera. Interestingly, we found obvious expansion of some subgroups in *E. onukii* (e.g., CSP3, CSP4, CSP8, and CSP9) and these CSPs were unevenly distributed over 4 of the 10 chromosomes, with enrichment on chromosome 1 (Fisher exact test, *P* value < 0.00001; Figure 2C). Most CSP genes were distributed in expanded clusters on chromosomes, likely through a series of gene duplication events (Figure 2C). Meanwhile, several CSPs were highly expressed across different life cycle stages (Figure S5A), implying an important role in the growth and development of *E. onukii.* Earlier studies on CSP functions [25, 26], coupled with our observations of conserved phylogeny in Hemiptera species and species-specific expansion of CSPs, indicate that CSPs are crucial for recognition of tea volatiles and location of potential host tea plants. We suggest that *E. onukii* requires many CSPs to specifically detect the complex molecular components of odors from different tea cultivars. Thus, our analyses highlighted directions for further experimental analysis of genes linked to host adaptation. Toward this goal, functional testing of CSPs might identify genes that were responsible for detection of specific tea cultivars by *E. onukii*.

**Figure 2.**
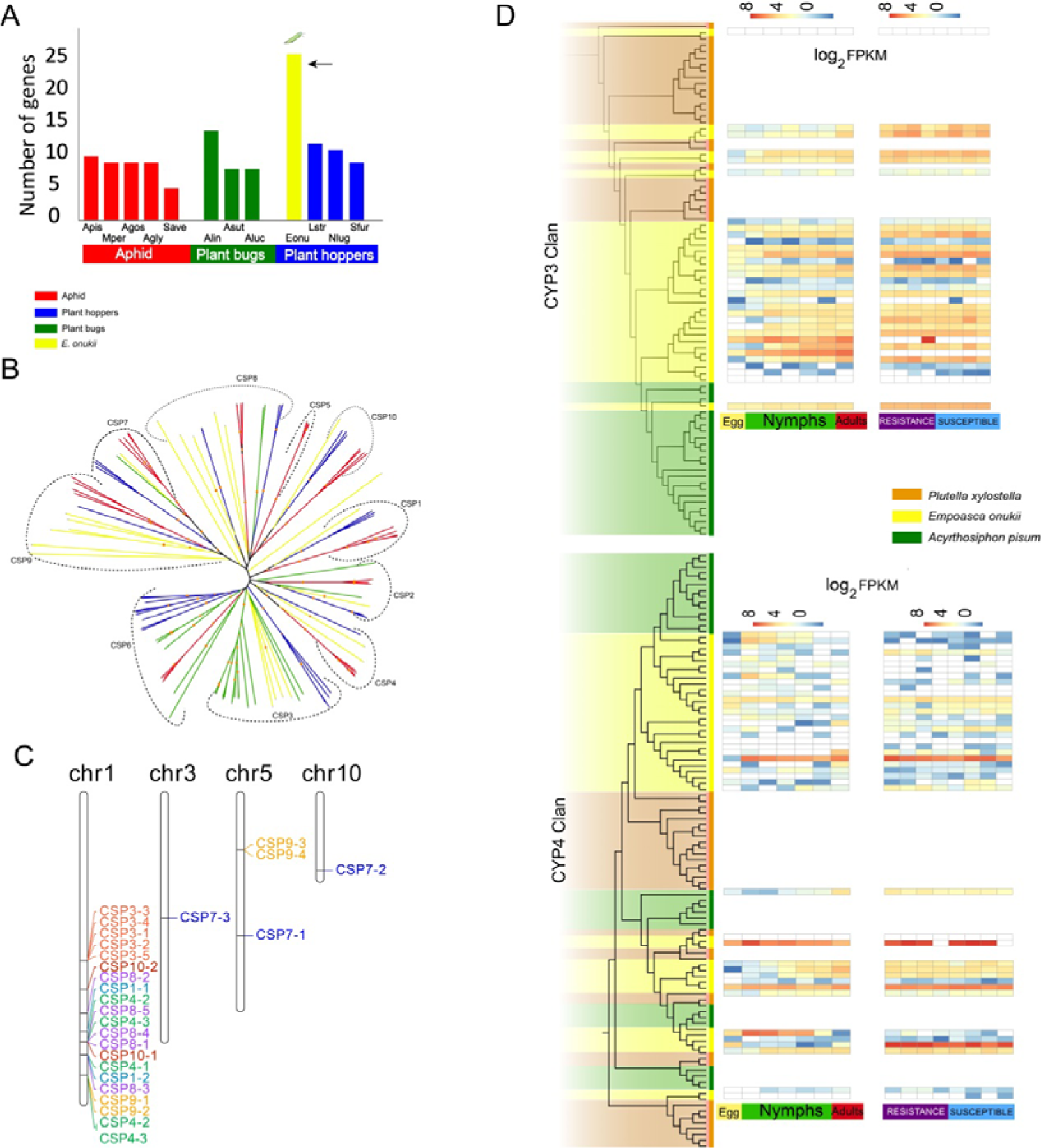
Expansion of gene families related to chemoreception and detoxification in *E. onukii* compared with other insect species. **(A)** Numerical comparison of the chemosensory proteins (CSPs) among aphids, plant bugs and hoppers. Aphid species include *Acyrthosiphon pisum* (Apis), *Myzus persicae* (Mper), *Aphis gossypii* (Agos), *Aphis glycines* (Agly) and *Sitobion avenae* (Save); Plant bugs include *Adelphocoris lineolatus* (Alin), *Adelphocoris suturalis* (Asut), *Apolygus lucorum* (Aluc); Hoppers includes *Empoasca onukii* (Eonu), *Laodelphax striatellus* (Lstr), *Nilaparvata lugens* (Nlug) and *Sogatella furcifera* (Sfur). **(B)** Phylogenetic relationships of CSPs in hoppers (*N. lugens*, *S. furcifera*, and *L*. *striatellus*), aphids (*A. pisum, M. persicae, A. gossypii, A. glycines*, and *S. avenae*), and plant bugs (*N. lugens, L. striatellus* and *S. furcifera*). Yellow branches represent CSP family genes in *E. onukii*. **(C)** Genomic expansion and unbalanced chromosomal distribution of CSPs in the *E. onukii* genome. **(D)** Phylogenetic relationships and expression profiling of detoxification-related proteins (CYP3 and CYP4) in plant hoppers, aphids and plant bugs. Expression profiling based on RNA-seq data were generated from all developmental stages (egg, 1st – 5th nymph instar and adult) and 8 *E. onukii* populations collected from different tea cultivars (four cultivars being resistant including LongJ, DeQ, JianD, JuY and four cultivars susceptible to *E. onukii* including ZhuS, LanT, BanZ and EnB).

Our investigation of the chemoreceptor-related genes showed relatively low numbers of ORs, IRs, GRs and OBPs in *E. onukii* (Table 3; Table S11; Figure S5-8). For example, we found that the number of OBPs in Hemiptera species was lower than other insect orders (Figure S5B), suggesting conservation of odorant molecular transport in Hemiptera [27]. Other chemoreceptor genes, including ORs, GRs and IRs, play important roles in local adaption by responding to chemical signals with neuronal activity [15]. The species-specific expansion OR clade of gene family was obvious in our analysis (Figure S6). Polyphagous insects (e.g., *P. americana*) possess more OR genes than monophagous insects (e.g., *E. onukii*) (Table 3; Table S11; Figure S6). This might have resulted from specific evolutionary adaption to food selection and detection since genetic diversity of ORs allows insects to bind to a greater range of ligands [28]. In addition, similar to *N. lugens, E. onukii* had a substantially lower number of GRs (Table 3, Figure S7). Earlier studies show a close relationship between GRs and insect herbivory, with lower number of GRs in specialists than in generalists [29, 30]. Another explanation may be that antennae of leafhoppers have a much simpler structure with fewer sensilla than those of planthoppers (e.g., *N. lugens*) and aphids. TGL also possessed fewer IRs (Table 3; Table S11; Figure S8), which mediate synaptic communication in insects and mediates responses to volatile chemicals in *D. melanogaster* [31, 32]. We believed that the numerical reduction in ORs, IRs, GRs and OBPs might be associated with the adaptive evolution to a monophagous diet of tea phloem sap, and the substantial expansion of CSPs might contribute to tea volatile perception in *E. onukii*.

*E. onukii* is believed to have experienced rapid evolution leading to insecticide resistance in natural populations [33]. Four classic gene families commonly associated with detoxification of xenobiotics and insecticides, including P450, COEs, GSTs and ABCs, were therefore investigated. We identified 103 cytochrome P450s, 29 ATP-binding cassette transporters (ABC transporters), 77 carboxylesterases (COEs), and 30 glutathione S-transferases (GSTs) (Table 3). Similar to other insecticide resistant pests [34], we found that the P450 gene family was expanded, mainly in CYP3 and CYP4 clans (Table 3; Table S12; Figure 2D; Figure S9A). Based on our RNA-seq data, 28 CYP3 and 38 CYP4 genes showed expression (FPKM > 1.0) with 20 in CYP3 and 16 in CYP4 being highly expressed during at least one developmental stage (FPKM > 10.0) (Figure 2D; Figure S9B). The results underlined their potential function of detoxifying the xenobiotics or insecticides in TGL.

We tracked the expression patterns of these genes in TGL samples collected from different tea cultivars, with four cultivars being resistant and four cultivars susceptible to TGL according to previous studies [35]. Results showed that 17 CYP3 and 12 CYP4 genes were highly expressed (Log_2_^FPKM^ > 10.0) but not differentiated in both resistant and susceptible tea cultivars (Figure 2D; Figure S9C). Thus, we speculated that CYP genes might be involved in metabolism of common xenobiotics, or their expression may be induced by insecticides. Indeed, previous studies have shown that CYP3s are involved in xenobiotic metabolism and insecticide resistance, with some family members being inducible by pesticides or plant secondary metabolites [36]. CYP4s are known to encode constitutive and inducible enzymes related to odorant and pheromone metabolism, and expression can be induced by xenobiotics [37].

### Niche under adaptive selection are related to metabolic regulation and detoxification

*E. onukii* samples were collected from four tea-growing regions around China: southwest region (SWR), south of the Yangtze River region (SYR), north of the Yangtze River region (NYR), and south China region (SER), and these samples were re-sequenced with a depth ranging from 20.8× to 30.7×at whole genome level (Table S13). After filtering the low-quality variants, we generated a genomic dataset containing 12,271,501 high-quality SNPs (Table S13) to estimate the genomic signatures of evolutionary adaptation for *E. onukii*, based on Tajima’s *D* with a 50-kb window size and a fixed step length of 10 kb. Totally 369 sliding windows, covering 18.45 Mb genomic sequences (Table S14) and containing 82 protein-coding genes (Table S15) were detected, including seven genes (Table S16) under purifying selection and 79 under balancing selection (**Figure 3**A).

**Figure 3.**
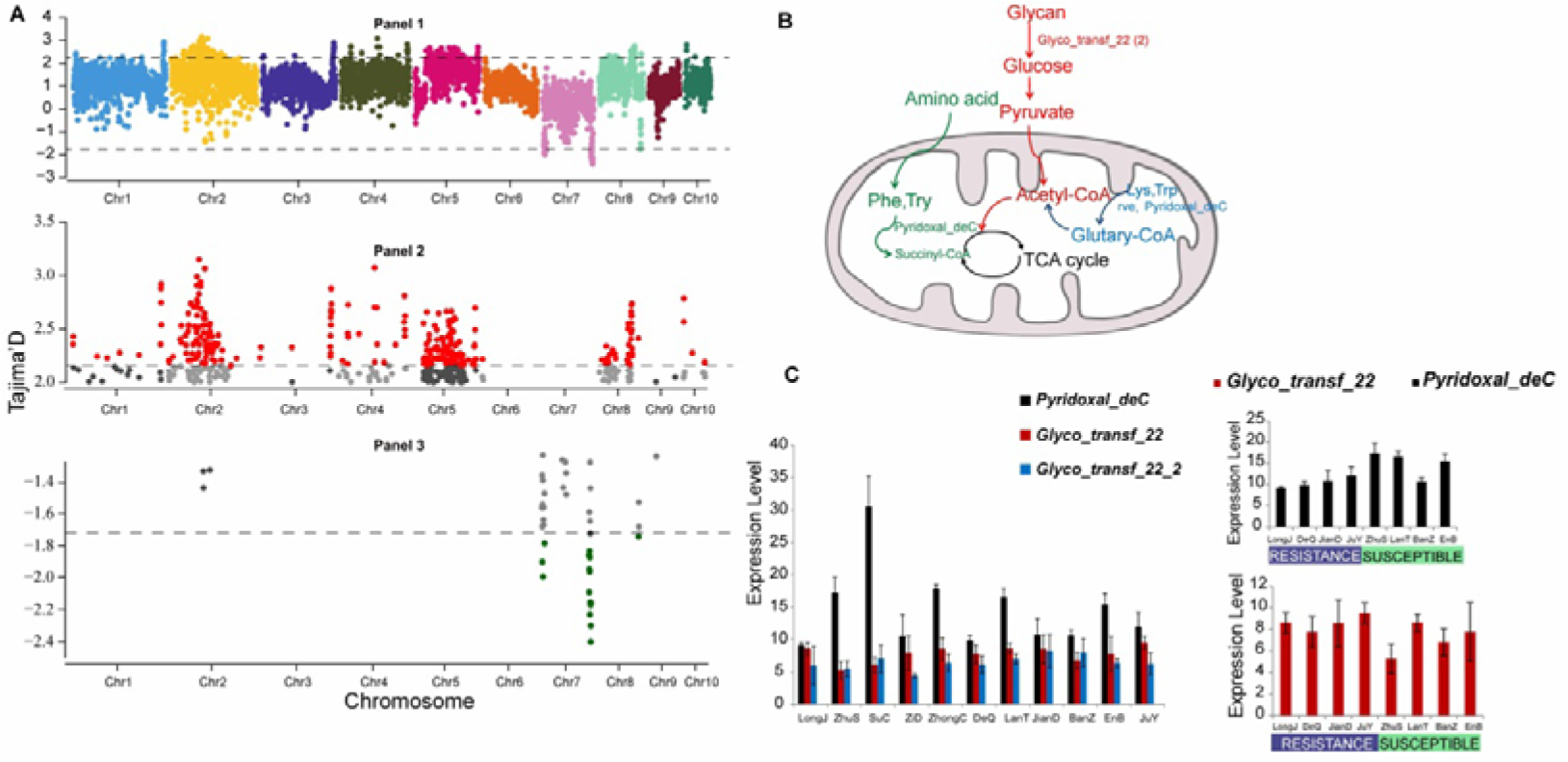
Genomic signatures of balancing selection. **(A)** Putative selection sweeps in populations of *E. onukii*. Tajima’s *D* value was calculated for each of the *E. onukii* populations. Mean values of Tajima’s *D* are shown in sliding windows of 50 kb with a step size of 10 kb. Regions with Tajima’s *D* values deviated significantly from 0 are marked with dotted lines in panel 1. Specifically, values of Tajima’s *D* significantly deviated from 0 are plotted in red (> 0) and green (< 0) respectively in panel 2 and panel 3. (B) Succinyl- and glutaryl-CoA pathways showing the regulatory role of lysine modifications in metabolism. (C) Expression patterns of the *E. onukii* genes under balancing selection, in 11 different tea cultivars. *Pyridoxal_deC* was detected to be significantly highly expressed in susceptible tea cultivars (*P* < 0.01, T-test), while *Glyco_transf_22* was significantly highly expressed in resistant tea cultivars (*P* < 0.05, T-test).

Purifying selection is important in shaping genomic diversity in natural populations and is essential to preserving biological functions at selection sites [38]. Almost all these genes are related to nervous system or visual functions. For instance, forkhead box protein P1 (*FOXP1*) is a transcription factor with a regulatory function in the central nervous system (CNS), and mutations in this gene have been linked to various neurodevelopmental diseases, including autism, cognitive abnormalities, intellectual disabilities and speech defects [39]. Coronin 6 is highly enriched at the adult neuromuscular junction and can regulate acetylcholine receptor (neurotransmitter receptor) clusters by modulating interactions between the actin cytoskeletal network and receptors [40]. In mice lacking *SZT2*, mTORC1 signaling is hyperactive in several tissues including neurons in the brain, and these components have been linked to neurological disease [17]. The nucleoredoxin-like 1 (*Nxnl1*) gene has two alternative splice isoforms: a rod-derived cone viability factor that functions in the retina [41]. Inositol hexakisphosphate, the nonvisual arrestin oligomerization and cellular localization are modulated by its binding [42]. In insects, detection of light changes, vibration, colors and semiochemicals, which have evolutionarily old sensory functions, are vital for behaviors including avoiding predation, food location and intraspecific communication. Thus, we speculated that these genes under purifying selection would be important for nervous or visual functions in *E. onukii*.

We found that 91.8% of the selected genes (79/86) were under balancing selection with positive Tajima’s *D* values that greatly deviated from zero, indicating that the populations of *E. onukii* maintained a high level of polymorphism. A much higher genetic diversity (π = 0.00804) was observed in those genomic regions under balancing selection compared to genome-wide diversity (*P* < 0.0001, T-test), suggesting a strong capacity for *E. onukii* to rapidly adapt to diverse habitats [43]. GO enrichment analysis showed that these genes were enriched in several biological processes including cell periphery, plasma membrane part, transmembrane transport, ion binding, anion binding, and nucleoside-triphosphatase activity (Figure S10; Table S17). KEGG enrichment analyses pointed to several pathways including those linked to metabolism, circadian rhythms, and immune system functions.

Based on KEGG analyses, lysine succinylation and glutarylation pathways were enriched (Figure 3B). Previous studies have reported that protein acetylation plays critical roles in cellular processes ranging from gene expression to metabolism [44]. Lysine succinylation is a recently identified post-translational modification (PTM) [45]. It is important for metabolism and detoxification in *B. mori* [45]. In our study, the apoptosis pathway was consistently enriched (Figure S11). Studies in Lepidoptera insects suggest that apoptosis plays a vital role in resistance to virus infection and some apoptosis-related proteins are known to be succinylated [40, 46, 47]. Here, we identified four key genes, including *rve, Pyridoxal_deC* and two *Glyco_transf_22*, which were functionally present in lysine succinylation and glutarylation pathways (Figure 3B).

Lysine succinylation is important for virus-infection resistance and detoxification in insects [44-46, 48, 49]. To examine whether the selected pathways in *E. onukii* had similar functions, we analyzed the expression patterns of four genes, *rve, Pyridoxal_deC* and two *Glyco_transf_22,* collected from 11 different tea cultivars including 4 cultivars (LongJ, DeQ, JianD, JuY) showing resistance to *E. onukii* and 4 cultivars (ZhuS, LanT, BanZ, EnB) showing susceptibility to *E. onukii*. Based on RNA-Seq analysis, two genes (*Pyridoxal_deC, Glyco_transf_22*) showed significantly high expression in both susceptible and resistant tea cultivars, but with different patterns (Figure 3C) (*P* < 0.05, T-test). *Pyridoxal_deC* and *Glyco_transf_22* were key genes in metabolic regulation of succinyl- and glutaryl-CoAs (Figure 3B). These results, together with the previously reported roles of succinylation and glutarylation in other insects [44-46, 49], indicated that genes under balancing selection could be involved in metabolic regulation and detoxification of *E. onukii*, possibly contributing to its success in adapting to a wide range of tea cultivars grown in ecologically diverse regions of China.

*E. onukii* is largely controlled using insecticides in China, leading to development of resistance to chemicals. The ATP-binding cassette (ABC) transporters are conserved across insects and have been implicated in insecticide resistance among pest species [50]. Based on our analyses, the ABC superfamily showed no expansion, and few orthologs were present in *E. onukii* compared to other insect species (Table 3). However, we identified four ABC transporter genes that showed signatures of balancing selection and thus maintaining high genetic variation within populations of *E. onukii*. Further analysis showed that these four genes belonged to three ABC subfamilies including ABCG, ABCB, and ABCA. Balancing selection favors defense proteins with functions in resistance, immunity and adaptations [51, 52]. However, the functions of these four genes have not been elucidated in leafhoppers or aphids. Studies in other animals or insects have shown that these subfamilies are closely related to drug or insecticide resistance [53, 54, 55]. A comparative analysis between susceptible and resistant strains of *A. aegypti* reports that the genes of ABC transporter G family are highly up-regulated [54]. Similar studies have also been carried out in *P. xylostella* and *L. striatellus*, showing that ABCA/B/G subfamilies are significantly over-expressed in the resistant strains [55, 56]. Based on ABC family functions in other insects, we hypothesized that *E. onukii* ABC genes might contribute to its adaptation to different tea cultivars. This hypothesis may be supported by a study of Cry1Ac resistance in *P. xylostella* [53]. We therefore investigated the expression patterns of the four genes using *E. onukii* samples collected from 11 different tea cultivars, as described above. These genes showed moderate expression levels across different developmental stages (Figure S12A) and samples of *E. onukii* from different tea cultivars (Figure S12B), suggesting that these ABC superfamily members could broadly contribute to adaptation to various tea cultivars and even possibly to chemical resistance. Previous studies also suggest that ABC transporters are not strictly specific to certain chemicals, implying that ABC transporters have a broad spectrum of chemical substrates and may act as a basis for cross-resistance of multiple chemicals [55].

Genomic regions under balancing selection are functionally important because of their high genetic diversity contributing to adaption to environmental change [43]. Based on our results, we hypothesized that balancing selection might have contributed to the high level of polymorphism in *E. onukii* populations, facilitating adaptation to diverse environments and tea cultivars.

### Evolutionary history is inconsistent between TGL and tea cultivars

We used high-quality SNPs obtained from the 54 *E. onukii* samples coming from the different tea-growing regions (**Figure 4**A) in China (Table S13) to profile their phylogeographical relationships. Phylogenetic analysis and network estimation of the *E. onukii* samples with *E. flavescens* and *Asymmetrasca* sp. as outgroups uncovered three geographically clustered groups (Groups I-III; Figure 4B and Figure S13). Group I contained 4 samples collected from Yunnan province, being the closest to the outgroups. Group II included 28 samples mainly collected from eastern China, including Shandong, Jiangsu, Zhejiang, and Anhui provinces. The remaining 22 samples (i.e., Group III) were mainly from 13 provinces of central and southern China (Figure 4B and **Table 4)**. These results were further supported by genetic structure analysis (*K* = 3) based on the Admixture model (Figure 4C and Figure S14) [57]. Three clustered groups of *E. onukii* samples (Figure 4B; Table 4) were inconsistent with the current division of the four tea-growing regions (Figure 4A) based on the tea growing history, geographical locations, and tea cultivars [58]. These results suggest a different evolutionary history of *E. onukii* among these regions. Our analyses of phylogenetic and genetic structure confirmed the genetic differences between group I (samples from Yunnan) and the other groups, as shown in a previous study based on microsatellites [5]. However, this previous study suggested four main genetic groups (*K* = 4) [5]. This may be because the present study collected much more samples around China (54 locations in 22 provinces) than the other one (22 19 locations in 13 provinces) and used a greater number of genetic markers (whole-genome SNPs vs. microsatellite markers). We observed that individuals from different groups were interspersed (Figures 4B and C), possibly reflecting gene flow across location, as observed in the previous study [5].

**Figure 4.**
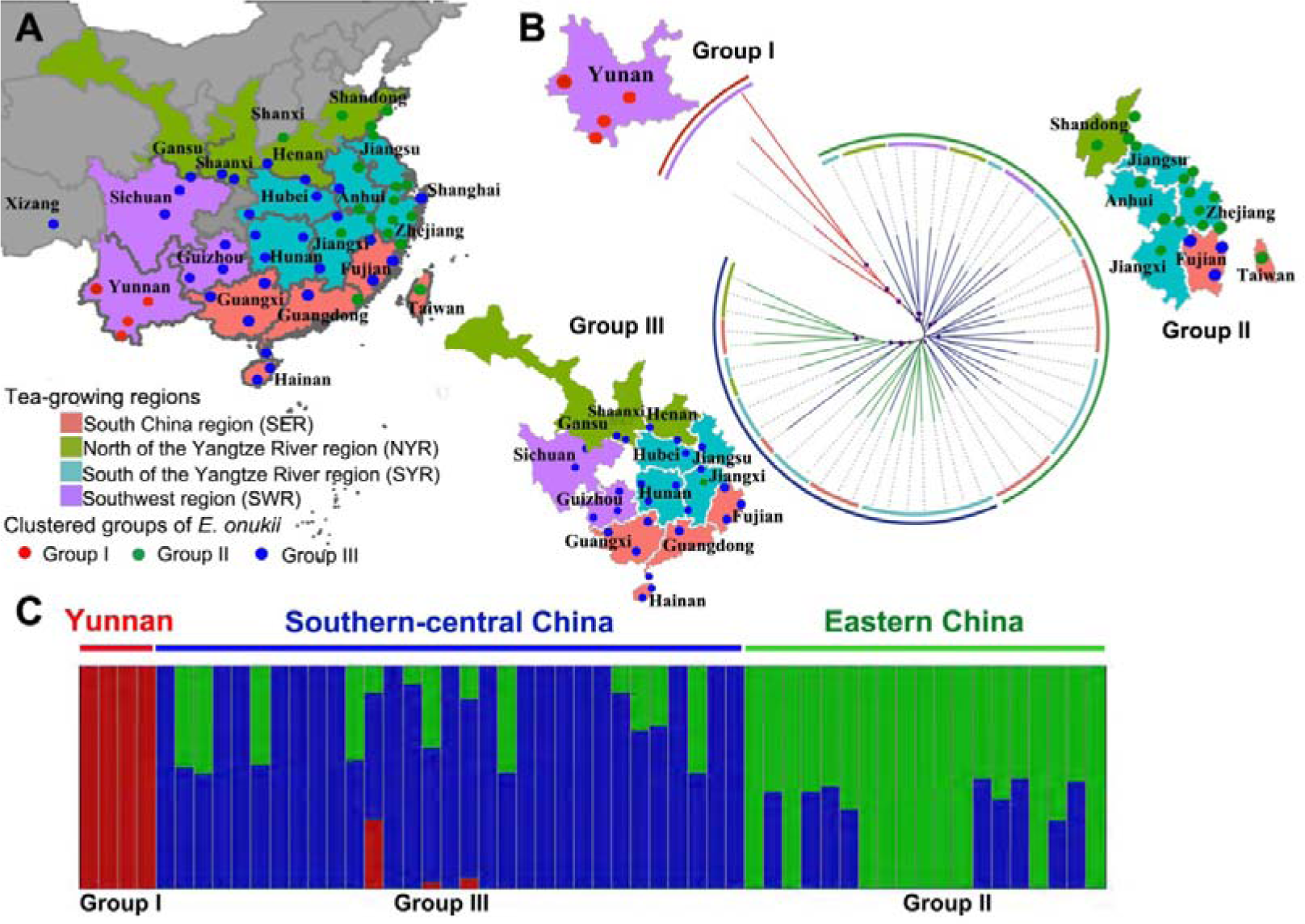
Phylogenetic relationship, population structure and expansion of *E. onukii*. **(A)** Geographical locations (sites) of 54 samples collected from four tea-growing regions around China: Southwest region (SWR), South of the Yangtze River region (SYR), North of the Yangtze River region (NYR), and South China region (SER). Dots with different colors represent different clustered groups. **(B)** Phylogenetic tree of the 54 *E. onukii* samples based on RAxML and SplitsTress. Branch lengths are not scaled. Different colors of inner circle represent 4 different tea-growing regions shown in (A). Colors of outer lane represent different *E. onukii* groups based on phylogenetic analysis. **(C)** Genetic structure and individual ancestry with colors in each column representing ancestry proportion over range of population sizes (*K* = 2-4, with an optimal *K* = 3).

To investigate the genetic divergence, we calculated the average pairwise diversity (π) within each of the clustered groups (Table 4). Comparably higher levels of genetic diversity (0.004662 and 0.004744) were observed in Groups II and III than in Group I (0.004062). The high genetic diversity of eastern China and southern-central China may be explained by a geographically wide range and ecologically diverse tea-growing conditions. Further, we found a higher genetic diversity in certain subgroups within the major tea-growing provinces of eastern China and southern-central China. The diversity (π = 0.0048) in Jiangsu and Zhejiang provinces was higher than the overall diversity of eastern China. and similarly, a higher diversity (π = 0.004939) was found in Anhui and Fujian provinces than in southern-central China, indicating the genetic diversity in these locations. We further analyzed the population differentiation (*F*_ST_) across different geographically clustered groups and showed a very low *F*_ST_value (0.005902) between Group II and III, indicating their genetically close relationship. In contrast, the *F*_ST_values between Group I and Group II or III were much higher (0.052321 in Group II *vs.* Group I; 0.043225 in Group III *vs.* Group I), suggesting that the samples from Yunnan province were genetically distant from the populations in eastern China and southern-central China.

A previous study reports that the structure of male genitalia varies among *E. onukii* locations from eastern China, southern-central China and Yunnan province [3]. Our population genetic analysis also showed genetic differences among these different groups (Figure 4). We speculated that some geographical barriers might have restricted gene flow leading to these differences. Yunnan is surrounded by mountains and rivers as a result of an uplift during the Quaternary and is isolated by the steep Hengduan Mountains. Unlike the clonal propagation of tea cultivars in other regions of China, tea cultivation in Yunnan has depended on seeds from early times [59]. This might have prevented the interbreeding of local *E. onukii* populations with populations from other regions. Similarly, the hilly region between Zhejiang and Fujian might separate *E. onukii* populations, leading to populations genetically different (Figure 4).

### Conclusions

In this project, we develop a high-quality chromosome-level genome with 92.7% BUSCO completeness for *E. onukii*, a species of crucial importance to a widely consumed crop linked to human health. Based on genomic profiling and comparison, we find complex patterns of genomic variation and expansion of gene families associated with evolutionary adaptation to chemosensory reception and xenobiotic detoxification. We identify genomic signatures of balancing selection to reveal the high genetic diversity of resistant genes, underlining their important roles in the adaptive evolution of *E. onukii*. Further, we analyze patterns of variation in genomic sequences from 54 samples and two outgroups, uncovering the population structure and evolutionary history of *E. onukii* across the four different tea-growing regions of China. This work will facilitate functional studies on the adaption of this pest to ecologically diverse habitats, and provide the genomic resources and genetic knowledge for development of sustainable pest management strategy.

## Materials and Methods

### Insect colony

*E. onukii* samples were collected in Fuzhou, Fujian province, southeastern China in July 2017 (on the tea cultivar of Huangdan), and then maintained on tea plants in the laboratory. The insectarium environment was set at 28 ± 1°C and 60 ± 5% RH with a photoperiod (light: dark = 12:12).

### Genome sequencing and assembly

Since the quality of *de novo* assembly is sensitive to genomic heterozygosity, genomic DNA of adults was extracted from insects after 12 generations of laboratory inbreeding. Chromosome-level assembly was performed using Nanopore (Oxford Nanopore Technologies, ONT) with chromatin conformation capture (Hi-C) technologies. The raw ONT reads were self-corrected using CANU version 1.7 [61] with parameter corOutCoverage = 100, and corrected reads were subject to two widely-used long-read assemblers, wtdbg2 [62] and SMARTdenovo (https://github.com/ruanjue/smartdenovo). These two assemblers applied the homopolyer compressed (HPC) k-mer indexing algorithm for sequence alignment and assembly, making the heterozygous regions prone to collapsing. To improve the contiguity of contig assemblies, we used Quickmerge [63] to reconcile wtdbg2 and SMARTdenovo assemblies. Each round of assemblies was inspected through evaluation of N50s based on assembled genome size as well as complete/duplicated BUSCO ratio (Table S3), showing 92.7% completeness and only 2.7% duplication. It also indicated that the redundant sequences were well-handled in our assembly. The total length of the final contig assembly for *E. onukii* genome was 599 Mb with a contig N50 size of 2.2 Mb. Illumina short reads were then used to polish the ONT assembled genome by Pilon [60] with the following parameters: --diploid --threads 6 --changes --tracks --fix bases --verbose --mindepth 4. Hi-C libraries were created from nymphs as previously described [34]. The original cross-linked fragments, also known as chimeric fragments, were then processed into paired-end sequencing libraries and sequenced on the Illumina HiSeq X-10 platform. Paired-end reads were uniquely mapped onto the draft assembly, then 3D-DNA pipeline [64] was recruited to correct any mis-joined contigs by detecting abrupt long-range contact patterns. The Hi-C corrected contigs were further linked into 10 pseudo-chromosomes using the ALLHiC pipeline [65]. In total, we generated 65 Gb data (∼109×) sequences for one cell by for Nanopore ONT and 37 Gb clean data (∼61×) of Illumina X-10 from polishing **(Table 1S)**.

### Official gene set annotation

Annotation of protein-coding genes was based on *ab initio* gene predictions, transcript evidence and homologous protein evidence, which was all implemented in the GEMOMA computational pipeline [66]. RNA-seq data were generated from every developmental stage (egg, 1st – 5th nymph instar and adult). Besides, multiple studies have shown that resistance to *E. onukii* varies with different tea cultivars [35, 67]. *E. onukii* samples were collected from 11 main tea cultivars in China, of which four were susceptible to *E. onukii* (ZhuS, LanT, BanZ and EnB), four were resistant (LongJ, DeQ, JianD, JuY), and three had unknown resistance status (SuC, ZiD and ZhongC) [35]. RNA-seq reads were first trimmed using the Trimmomatic program [68] and then mapped to the reference genome using HiSAT2 [69]. During homolog-based prediction, the protein sequences of *Drosophila melanogaster*, *Apis mellifera*, *Myzus persicae*, *Acyrthosiphon pisum*, *Tribolium castaneum*, and *Bombyx mori* were downloaded and aligned to the reference assembly using TBLASTN with e-value 1e^-5^, and the resulting alignment files were subject to GEMOMA annotation.

### Orthology and phylogenomics

A total of 19 representative insect species including *E. onukii* were collected for orthology and phylogenetic analyses (**Figure 1**). A phylogenetic tree based on a concatenated sequence alignment of the single-copy gene families from *E. onukii* and other insect species was constructed. We identified 5,736 single copy genes in these insect genomes using OrthoFinder (version 2.0.0) [70] and performed multiple alignments of the single copy genes from the selected genomes using MAFFT v7.299b [71]. Based on a concatenated sequence alignment, a phylogenetic tree was constructed using RAxML software and the PROTGAMMALGX model [18]. Divergence times of the selected insect species were calculated by PAMLv4.8a mcmcTREE [72]. The Markov chain Monte Carlo (MCMC) was run for 1,000,000 iterations using a sample frequency of 100 after a burn-in of 2,000 iterations, with the other parameters set as defaults. The following constraints were used for time calibrations: *D. melanogaster* and *A. mellifera* divergence time (42.8-83.4 million years ago) [73]. FigTree v1.44 was used to visualize the phylogenetic tree. Gene family expansion and contraction analyses were performed using Café v4.0.1 [74].

### Gene families

Some gene families with functional importance were selected for manual annotation based on the high-quality assembly. Most gene families were annotated using known models from previously annotated genomes including *D. melanogaster*, *A. mellifera, M. persicae, A. pisum, C. lectularius* and *B. mori*. Some gene families, which were difficult to identify from automated predictions, were identified based on iterative searching. In brief, BLASTP searches for Hemiptera homologs used queries to search the genomic loci for significant hits (e < 10^-3^).

Further, we recruited hidden Markov models (HMMs) to identify certain domains for these selected gene families based on pfam_scan [75]. Multiple sequence alignments of the selected gene families were obtained with MUSCLE [76] and corrected manually. Phylogenetic analysis was conducted using ML and NJ models, and implemented in MEGA7 for 500 bootstraps [77].

### Differential gene expression

RNA-seq data were generated from seven developmental stages (egg, 1^st^ – 5^th^nymph instars and adult) and the 11 populations of *E. onukii* collected from different tea cultivars (described in Official gene set annotation section). The RNA-seq reads were trimmed using the Trimmomatic program [68] and mapped back to gene models using bowtie [78]. FPKM was calculated based on the RSEM program [79] implanted in Trinity software [80]. Significantly differentially expressed genes were detected with a cutoff (*P* < 0.05 and log_2_^(changefold)^ > |1|) [81].

### Sample collection, resequencing and SNP calling

Individuals of *E. onukii* (between 100 and 120) were collected from 54 plantations distributed in four tea-growing regions of China: Southwest region (SWR), South of the Yangtze River region (SYR), North of the Yangtze River region (NYR), and South China region (SER) (**Table S13**). *W*e also collected two samples, *E. flavescens* (collected from Canada in vineyards) and *Asymmetrasca* sp. (collected from Africa), as outgroups (Table S13). For DNA extraction, 50 to 100 individuals were mixed. Genomic DNA extraction, library construction and amplification were performed following standard protocols (Supplemental Notes). All samples were sequenced using the Illumina X-10 platform with a paired-end read length of 150 bp. The GATK toolkit (version: V 3.5-0-g36282e4) [82] and samtools/bcftools [83] were used to detect variants and SNPs following a series of filtering steps as detailed in a Supplemental Note.

### Maximum-likelihood tree inference

The phylogenetic tree was built based on SNPs of single-copy genes. The heterozygous and homozygous SNPs were included in the construction of ML tree. For the heterozygous SNPs, the major alleles that had more reads supported than the minor alleles were retained for further analysis. These SNPs were converted to phylip and aligned in fasta format. The ML (maximum likelihood) tree was constructed using IQ-Tree with a self-estimated best substitution model [84].

### Admixture analysis

Ancestral population stratification among the re-sequenced TGL populations was inferred using Admixture software [85]. We estimated the optimal ancestral population structure using ancestral population sizes *K* = 1 – 4 and estimated parameter standard errors based on bootstrapping of 2000.

### Diversity statistics

VCFtools v 0.1.3 [86] was used to calculate population diversity statistics. Genetic differentiation (*F*_ST_) and the average pairwise diversity index (π) were estimated based on a sliding window analysis with 100 kb window size and 50 kb step size.

### Scanning loci under selective sweeps

To identify candidate genes responsible for reciprocal selection in the TGL populations, we performed the Tajima’s *D* test to identify selective sweeps. Loci with Tajima’s *D* that greatly deviated from 0 proved to be a selection niche in the genome. The Tajima’s D statistics were calculated using VCFtools program with a 50 kb window size and 10 kb step size. A negative Tajima’s *D* indicates population size expansion and/or purification selection. A significantly positive Tajima’s *D* signifies low levels of low and high frequency polymorphisms, indicating a decrease in population size and/or balancing selection [87]. We used the empirical 5% windows to indicate the significance. The lowest 5% windows were considered as purifying selection and the highest 5% were considered as balancing selection.

Based on the annotation of our high-quality genome, candidate genes were identified using our outliers. GO annotation was conducted using Blast2GO [88] and the KEGG pathway analysis was performed using OmicShare tools (www.omicshare.com/tools).

## Data availability

The genome sequences and re-sequencing reads have been deposited in NCBI with accession number of PRJNA731240 and GSA database (https://ngdc.cncb.ac.cn/search/?dbId=gsa&q=Empoasca) with the accession number of GWHBAZN00000000. Reads for RNA-seq was deposited in GSA database with the accession number of PRJCA005189. The mitochondrial sequence reported in this paper have been deposited in the Genome Warehouse in National Genomics Data Center with the accession number GWHBFSP00000000 that is publicly accessible at https://ngdc.cncb.ac.cn/gwh.

## CRediT author statement

**Qian Zhao:** Investigation, Methodology, Formal analysis, Visualization, Writing – original draft, Writing – review & editing. **Longqing Shi:** Resources, Methodology, Writing – original draft. **Weiyi He:** Methodology, Writing - review & editing. **Jinyu Li:** Resources, Formal analysis. **Shijun You:** Resources, Data curation. **Shuai Chen:** Formal analysis. **Jing Lin:** Visualization, Formal analysis. **Yibin Wang:** Formal analysis. **Liwen Zhang:** Visualization. **Guang Yang:** Resources, Writing - review & editing. **Liette Vasseur:** Resources, Writing - review & editing. **Minsheng You:** Conceptualization, Methodology, Resources, Writing - original draft, Writing - review & editing, Supervision. All authors read and approved the final manuscript.

## Acknowledgements

This work was supported by The National Key R & D Program of China (2019YFD1002100), Fujian Agriculture and Forestry University Construction Project for Technological Innovation and Service System of Tea Industry Chain (K1520005A03) and Key International Science and Technology cooperation Project of China (2016YFE0102100). We thank Mr. Haifang He, and Fasheng Huang for their kind assistance in collection of the insect samples.

## Supplementary material

Title for supplementary file: Supplemental_Notes.docx

## Figure legends

**Figure S1. Damages of *E.onukii* in modern tea plantation in China.** (A) Life cycle of *E. onukii*. (B) Damages caused by *E.onukii*. (C) Developmental stages of *E. onukii*.

**Figure S2. Genome assembly of *E. onukii*. (A) Chromatin interactions with 150 kb resolution in *E. onukii.* (B) Mitochondrial genome of *E. onukii.*** Inner circle represents the GC content while outer circle represents genes located on mitochondrion.

**Figure S3. (A) Assessment of the assembly using LTR Assembly Index (LAI). (B) Correlation analysis between repeat content and the genome size.**

**Figure S4. Gene family expansions and contractions in the *E. onukii* compared with other insects.** Numbers for expanded (green) and contracted (red) gene families are shown on branches.

**Figure S5. Expression profile of genes involved in chemoreception and phylogenetic analysis of odorant-binding proteins (OBPs).** (A) Expression of 5 chemoreception gene families. (B) Neighbor-joining method involving 84 protein sequences was used to construct the tree. Different colors represent different species: red represents *Ap. Glycines* (Agly); green represents *A. pisum* (Apis); purple represents *E. onukii* (Em) while black represents *D. melanogaster*.

**Figure S6. Phylogenetic analysis of olfactory receptors (ORs).** Neighbor-joining method involving 216 protein sequences was used to construct the tree. Different colors represent different species: red represents *Ap. Glycines* (Agly); green represents *A. pisum* (Apis); yellow represents *E. onukii* (Em) while blue represents *D. melanogaster*.

**Figure S7. Phylogenetic analysis of gustatory receptors (GRs).** Neighbor-joining method involving 219 protein sequences was used to construct the tree. Colors corresponded to Supplemental Figure 6.

**Figure S8. Phylogenetic analysis of ionotropic receptors (IRs).** Neighbor-joining method involving 125 protein sequences was used to construct the tree. Colors corresponded to Supplemental Figure 6.

**Figure S9. Phylogenetic analysis of P450 gene family.** Neighbor-joining method involving 259 protein sequences was used to construct the tree. Colors represented different species listed in the Figure.

**Figure S10. GO enrichment of the genes under balancing selection in *E. onukii*. Only top 20 were listed in the figures.**

**Figure S11. Apoptosis pathway was enriched for the genes under balancing selection in *E. onukii*. Genes under selection were marked in red.**

**Figure S12. Expression patterns of the ABC genes under balancing selection.** (A) Expression patterns of different developmental stages; (B) Expression patterns when living on resistant and susceptible tea cultivars.

**Figure S13. Phylogenetic tree and network estimation by RAxML and SplitsTress.** Three populations according to the geographic regions that most of the individuals located: group I was Yunan (YN), Eastern China (group II); and central and southern of China (group III) were listed in Red, Green, Blue respectively (right figure) with presence of the outgroups (left figure). Different 4 tea regions of China were listed in different colors represented in Figure 3.

**Figure S14. Admixture analysis of 55 TGLs accessions (*k* = 2-4). IDs were represented in Supplemental Table 12**.

## Table legends

Table S1. Statistics of genomic sequencing data of Empoasca onukii

Table S2. Statistics of Hi-C mapping

Table S3. Statistics of contig level assembly of E. onukii

Table S4. BUSCO analysis of genome assembly of E. onukii

Table S5. BUSCO analysis of annotation completeness

Table S6. The statistics of different Hemiptera species assembly

Table S7. Assessment of genome consistency based on NGS (Illumina) reads

Table S8. Statistics of TEs in E. onukii genome

Table S9. GO over-representation of gene families expanded on *E.onukii* branch

Table S10. Gene family contraction analysis on *E.onukii* branch

Table S11. Chemosensory related gene families in E. onukii

Table S12. List of gene families involved in detoxification

Table S13. Geographic distributions of the collected samples around China

Table S14. Genomic regions under selection

Table S15. Genes under selection

Table S16. Functional analysis of genes under purifying selection

Table S17. GO terms for the genes under balancing selection

## References

1. Fu JY, Han BY and Xiao Q. Mitochondrial COI and 16sRNA Evidence for a Single Species Hypothesis of *E. vitis*, J. formosana and E. onukii in East Asia. PLoS One 2014; 9(12): p. e115259.

2. Chen LL, Yuan P, Pozsgai G, Chen P, Zhu H and You MS. The impact of cover crops on the predatory mite *Anystis baccarum* (Acari, Anystidae) and the leafhopper pest *Empoasca onukii* (Hemiptera, Cicadellidae) in a tea plantation. Pest Manag Sci 2019; 75(12): p. 3371–3380.

3. Qin D, Zhang L, Xiao Q, Dietrich C and Matsumura M. Clarification of the Identity of the Tea Green Leafhopper Based on Morphological Comparison between Chinese and Japanese Specimens. PLoS One 2015; 10(9): p. e0139202.

4. Lv WM, Chen X and Luo QR. Research on occurrence and control of *Empoasca flavescens*. Journal of Tea Science 1964: p. 45–55.

5. Zhang L, Wang F, Qiao L, Dietrich CH, Matsumura M and Qin D. Population structure and genetic differentiation of tea green leafhopper, *Empoasca (Matsumurasca) onukii*, in China based on microsatellite markers. Sci Rep 2019; 9(1): p. 1202.

6. Xiao Z, Huang X, Zang Z and Yang H. Spatio-temporal variation and the driving forces of tea production in China over the last 30 years. J. Geogr. Sci 2018; 28: p. 275–290.

7. Panfilio KA, Vargas Jentzsch IM, Benoit JB, Erezyilmaz D, Suzuki Y, Colella S, et al. Molecular evolutionary trends and feeding ecology diversification in the Hemiptera, anchored by the milkweed bug genome. Genome Biol 2019; 20(1): p. 64.

8. Jin S, Sun X, Chen Z and Xiao B. Resistance of the tea green leafhopper to different tea plant varieties. Sci. Agric. Sin 2012; 45(2): p. 255–265.

9. Rosenfeld JA, Reeves D, Brugler MR, Narechania A, Simon S, Durrett R, et al. Genome assembly and geospatial phylogenomics of the bed bug *Cimex lectularius*. Nat Commun 2016; 7: p. 10164.

10. J. A. Wenger, B. J. Cassone, F. Legeai, J. S. Johnston, R. Bansal, A. D. Yates, et al. Whole genome sequence of the soybean aphid, *Aphis glycines*. Insect Biochem Mol Biol 2017.

11. Ou S, Chen J and Jiang N. Assessing genome assembly quality using the LTR Assembly Index (LAI). Nucleic Acids Res 2018; 46(21): p. e126.

12. Nicholson SJ, Nickerson ML, Dean M, Song Y, Hoyt PR, Rhee H, et al. The genome of *Diuraphis noxia*, a global aphid pest of small grains. BMC Genomics 2015; 16: p. 429.

13. Li Y, Park H, Smith TE and Moran NA. Gene Family Evolution in the Pea Aphid Based on Chromosome-Level Genome Assembly. Mol Biol Evol 2019; 36(10): p. 2143–2156.

14. Ye YX, Zhang HH, Li DT, Zhuo JC, Shen Y, Hu QL, et al. Chromosome-level assembly of the brown planthopper genome with a characterized Y chromosome. Mol Ecol Resour 2021; 21(4): p. 1287–1298.

15. Xue J, Zhou X, Zhang CX, Yu LL, Fan HW, Wang Z, et al. Genomes of the rice pest brown planthopper and its endosymbionts reveal complex complementary contributions for host adaptation. Genome Biol 2014; 15(12): p. 521.

16. Kapusta A, Suh A and Feschotte C. Dynamics of genome size evolution in birds and mammals. Proc Natl Acad Sci U S A 2017; 114(8): p. E1460–E1469.

17. Wolfson RL, Chantranupong L, Wyant GA, Gu X, Orozco JM, Shen K, et al. KICSTOR recruits GATOR1 to the lysosome and is necessary for nutrients to regulate mTORC1. Nature 2017; 543(7645): p. 438–442.

18. Stamatakis A. RAxML version 8: a tool for phylogenetic analysis and post-analysis of large phylogenies. Bioinformatics 2014; 30(9): p. 1312–3.

19. F. Cui, M. X. Li, H. J. Chang, Y. Mao, H. Y. Zhang, L. X. Lu, et al. Carboxylesterase-mediated insecticide resistance: Quantitative increase induces broader metabolic resistance than qualitative change. Pestic Biochem Physiol 2015; 121: p. 88–96.

20. S. Cheng, J. Ashley, J. D. Kurleto, M. Lobb-Rabe, Y. J. Park, R. A. Carrillo, et al. Molecular basis of synaptic specificity by immunoglobulin superfamily receptors in Drosophila. Elife 2019; 8.

21. C. Xu, E. Theisen, R. Maloney, J. Peng, I. Santiago, C. Yapp, et al. Control of Synaptic Specificity by Establishing a Relative Preference for Synaptic Partners. Neuron 2020; 106(2): p. 355.

22. Dahanukar A, Hallem EA and Carlson JR. Insect chemoreception. Curr Opin Neurobiol 2005; 15(4): p. 423–30.

23. Bargmann CI. Comparative chemosensation from receptors to ecology. Nature 2006; 444(7117): p. 295-301.

24. Wang Q, Zhou JJ, Liu JT, Huang GZ, Xu WY, Zhang Q, et al. Integrative transcriptomic and genomic analysis of odorant binding proteins and chemosensory proteins in aphids. Insect Mol Biol 2019; 28(1): p. 1–22.

25. Youn YN. Electroantennogram responses of *Nilaparvata lugens* (Homoptera: Delphacidae) to plant volatile compounds. J Econ Entomol 2002; 95(2): p. 269–77.

26. He P, Zhang J, Liu NY, Zhang YN, Yang K and Dong SL. Distinct expression profiles and different functions of odorant binding proteins in *Nilaparvata lugens* Stal. PLoS One 2011; 6(12): p. e28921.

27. Xue J, Zhou X, Zhang XC, Yu LL, Fan HW, Wang Z, et al. Genomes of the rice pest brown planthopper and its endosymbionts reveal complex complementary contributions for host adaptation. Genome Biol 2014; 15: p. 521.

28. Robertson HM, Robertson ECN, Walden KKO, Enders LS and Miller NJ. The chemoreceptors and odorant binding proteins of the soybean and pea aphids. Insect Biochem Mol Biol 2019; 105: p. 69–78.

29. Pearce SL, Clarke DF, East PD, Elfekih S, Gordon KHJ, Jermiin LS, et al. Genomic innovations, transcriptional plasticity and gene loss underlying the evolution and divergence of two highly polyphagous and invasive Helicoverpa pest species. BMC Biol 2017; 15(1): p. 63.

30. McBride CS. Rapid evolution of smell and taste receptor genes during host specialization i n *Drosophila sechellia*. Proc Natl Acad Sci U S A 2007; 104(12): p. 4996–5001.

31. Benton R, Vannice KS, Gomez-Diaz C and Vosshall LB. Variant ionotropic glutamate receptors as chemosensory receptors in Drosophila. Cell 2009; 136(1): p. 149–62.

32. Chen C, Buhl E, Xu M, Croset V, Rees JS, Lilley KS, et al. Drosophila Ionotropic Receptor 25a mediates circadian clock resetting by temperature. Nature 2015; 527(7579): p. 516–20.

33. Wei Q, Yu HY, Niu CD, Yao R, Wu SF, Chen Z, et al. Comparison of Insecticide Susceptibilities of *Empoasca vitis* (Hemiptera: Cicadellidae) from Three Main Tea-Growing Regions in China. J Econ Entomol 2015; 108(3): p. 1251–9.

34. Wan F, Yin C, Tang R, Chen M, Wu Q, Huang C, et al. A chromosome-level genome assembly of *Cydia pomonella* provides insights into chemical ecology and insecticide resistance. Nat Commun 2019; 10(1): p. 4237.

35. Jin S, Sun XL, Chen Z and Xiao B. Resistance of different tea cultivars to *Emposca vitis* gOTHE. Sci. Agric. Sin 2012; 45(2): p. 255–265.

36. Feyereisen R. Evolution of insect P450. Biochem Soc Trans 2006; 34(Pt 6): p. 1252–5.

37. Simpson AE. The cytochrome P450 4 (CYP4) family. Gen Pharmacol 1997; 28(3): p. 351–9.

38. ] Cvijovic I, Good BH and Desai MM. The Effect of Strong Purifying Selection on Genetic Diversity. Genetics 2018; 209(4): p. 1235-1278.

39. Braccioli L, Nijboer CH and Coffer PJ. Forkhead box protein P1, a key player in neuronal development? Neural Regen Res 2018; 13(5): p. 801–802.

40. Chen Y, Ip FC, Shi L, Zhang Z, Tang H, Ng YP, et al. Coronin 6 regulates acetylcholine receptor clustering through modulating receptor anchorage to actin cytoskeleton. J Neurosci 2014; 34(7): p. 2413–21.

41. Byrne LC, Dalkara D, Luna G, Fisher SK, Clerin E, Sahel JA, et al. Viral-mediated RdCVF and RdCVFL expression protects cone and rod photoreceptors in retinal degeneration. J Clin Invest 2015; 125(1): p. 105–16.

42. Milano SK, Kim YM, Stefano FP, Benovic JL and Brenner C. Nonvisual arrestin oligomerization and cellular localization are regulated by inositol hexakisphosphate binding. J Biol Chem 2006; 281(14): p. 9812–23.

43. Wu J, Wang Y, Xu J, Korban SS, Fei Z, Tao S, et al. Diversification and independent domestication of Asian and European pears. Genome Biol 2018; 19(1): p. 77.

44. Hirschey MD and Zhao Y. Metabolic Regulation by Lysine Malonylation, Succinylation, and Glutarylation. Mol Cell Proteomics 2015; 14(9): p. 2308–15.

45. Chen J, Li F, Liu Y, Shen W, Du X, He L, et al. Systematic identification of mitochondrial lysine succinylome in silkworm (*Bombyx mori*) midgut during the larval gluttonous stage. J Proteomics 2018; 174: p. 61–70.

46. Cheng Y, Wang XY, Hu H, Killiny N and Xu JP. A hypothetical model of crossing *Bombyx mori* nucleopolyhedrovirus through its host midgut physical barrier. PLoS One 2014; 9(12): p. e115032.

47. Wang XY, Yu HZ, Xu JP, Zhang SZ, Yu D, Liu MH, et al. Comparative Subcellular Proteomics Analysis of Susceptible and Near-isogenic Resistant *Bombyx mori* (Lepidoptera) Larval Midgut Response to BmNPV infection. Sci Rep 2017; 7: p. 45690.

48. Gu Z, Zhou Y, Xie Y, Li F, Ma L, Sun S, et al. The adverse effects of phoxim exposure in the midgut of silkworm, *Bombyx mori*. Chemosphere 2014; 96: p. 33–8.

49. Sagisaka A, Fujita K, Nakamura Y, Ishibashi J, Noda H, Imanishi S, et al. Genome-wide analysis of host gene expression in the silkworm cells infected with *Bombyx mori* nucleopolyhedrovirus. Virus Res 2010; 147(2): p. 166–75.

50. Rosner J and Merzendorfer H. Transcriptional plasticity of different ABC transporter genes from *Tribolium castaneum* contributes to diflubenzuron resistance. Insect Biochem Mol Biol 2020; 116: p. 103282.

51. Koenig D, Hagmann J, Li R, Bemm F, Slotte T, Neuffer B, et al. Long-term balancing selection drives evolution of immunity genes in Capsella. Elife 2019; 8.

52. Van der Hoorn RA, De Wit PJ and Joosten MH. Balancing selection favors guarding resistance proteins. Trends Plant Sci 2002; 7(2): p. 67–71.

53. Ocelotl J, Sanchez J, Gomez I, Tabashnik BE, Bravo A and Soberon M. ABCC2 is associated with Bacillus thuringiensis Cry1Ac toxin oligomerization and membrane insertion in diamondback moth. Sci Rep 2017; 7(1): p. 2386.

54. Lien NTK, Ngoc NTH, Lan NN, Hien NT, Tung NV, Ngan NTT, et al. Transcriptome Sequencing and Analysis of Changes Associated with Insecticide Resistance in the Dengue Mosquito (*Aedes aegypti*) in Vietnam. Am J Trop Med Hyg 2019; 100(5): p. 1240–1248.

55. Sun H, Pu J, Chen F, Wang J and Han Z. Multiple ATP-binding cassette transporters are involved in insecticide resistance in the small brown planthopper, *Laodelphax striatellus*. Insect Mol Biol 2017; 26(3): p. 343–355.

56. You M, Yue Z, He W, Yang X, Yang G, Xie M, et al. A heterozygous moth genome provides insights into herbivory and detoxification. Nat Genet 2013; 45(2): p. 220–5.

57. Pritchard JK, Stephens M and Donnelly P. Inference of population structure using multilocus genotype data. Genetics 2000; 155(2): p. 945–59.

58. Zhang WJ, Rong J, Wei CL, Gao LP and Chen JK. Domestication origin and spread of cultivated tea plants. Biodiversity Science 2018; 26(4): p. 357–372.

59. Preparation committee, Records of tea varieties in China. 2001, Shanghai: Shanghai Scientific & Technical Publishers.

60. Walker BJ, Abeel T, Shea T, Priest M, Abouelliel A, Sakthikumar S, et al. Pilon: an integrated tool for comprehensive microbial variant detection and genome assembly improvement. PLoS One 2014; 9(11): p. e112963.

61. Koren S, Walenz BP, Berlin K, Miller JR, Bergman NH and Phillippy AM. Canu: scalable and accurate long-read assembly via adaptive k-mer weighting and repeat separation. Genome Res 2017; 27(5): p. 722–736.

62. Ruan J and Li H. Fast and accurate long-read assembly with wtdbg2. Nat Methods 2020; 17(2): p. 155–158.

63. Chakraborty M, Baldwin-Brown JG, Long AD and Emerson JJ. Contiguous and accurate de novo assembly of metazoan genomes with modest long read coverage. Nucleic Acids Res 2016; 44(19): p. e147.

64. Dudchenko O, Batra SS, Omer AD, Nyquist SK, Hoeger M, Durand NC, et al. De novo assembly of the Aedes aegypti genome using Hi-C yields chromosome-length scaffolds. Science 2017; 356(6333): p. 92-95.

65. Zhang X, Zhang S, Zhao Q, Ming R and Tang H. Assembly of allele-aware, chromosomal-scale autopolyploid genomes based on Hi-C data. Nat Plants 2019; 5(8): p. 833–845.

66. ] Keilwagen J, Hartung F, Paulini M, Twardziok SO and Grau J. Combining RNA-seq data and homology-based gene prediction for plants, animals and fungi. BMC Bioinformatics 2018; 19(1): p. 189.

67. Miao J, Han BY and Zhang QH. Probing behavior of *Empoasca vitis* (Homoptera: Cicadellidae) on resistant and susceptible cultivars of tea plants. J Insect Sci 2014; 14.

68. Bolger AM, Lohse M and Usadel B. Trimmomatic: a flexible trimmer for Illumina sequence data. Bioinformatics 2014; 30(15): p. 2114–20.

69. Kim D, Paggi JM, Park C, Bennett C and Salzberg SL. Graph-based genome alignment and genotyping with HISAT2 and HISAT-genotype. Nat Biotechnol 2019; 37(8): p. 907–915.

70. Emms DM and Kelly S. OrthoFinder: phylogenetic orthology inference for comparative genomics. Genome Biol 2019; 20(1): p. 238.

71. Nakamura T, Yamada KD, Tomii K and Katoh K. Parallelization of MAFFT for large-scale multiple sequence alignments. Bioinformatics 2018; 34(14): p. 2490–2492.

72. Yang Z. PAML: a program package for phylogenetic analysis by maximum likelihood. Comput Appl Biosci 1997; 13(5): p. 555–6.

73. Xiao JH, Yue Z, Jia LY, Yang XH, Niu LH, Wang Z, et al. Obligate mutualism within a host drives the extreme specialization of a fig wasp genome. Genome Biol 2013; 14(12): p. R141.

74. De-Bie T, Cristianini N, Demuth JP and Hahn MW. CAFE: a computational tool for the study of gene family evolution. Bioinformatics 2006; 22(10): p. 1269–71.

75. Eddy SR. Profile hidden Markov models. Bioinformatics 1998; 14(9): p. 755–63.

76. Edgar RC. MUSCLE: multiple sequence alignment with high accuracy and high throughput. Nucleic Acids Res 2004; 32(5): p. 1792–7.

77. Hall BG. Building phylogenetic trees from molecular data with MEGA. Mol Biol Evol 2013; 30(5): p. 1229–35.

78. Langmead B, Trapnell C, Pop M and Salzberg SL. Ultrafast and memory-efficient alignment of short DNA sequences to the human genome. Genome Biol 2009; 10(3): p. R25.

79. Li B and Dewey CN. RSEM: accurate transcript quantification from RNA-Seq data with or without a reference genome. BMC Bioinformatics 2011; 12: p. 323.

80. Haas BJ, Papanicolaou A, Yassour M, Grabherr M, Blood PD, Bowden J, et al. De novo transcript sequence reconstruction from RNA-seq using the Trinity platform for reference generation and analysis. Nat Protoc 2013; 8(8): p. 1494–512.

81. Montgomery SH and Mank JE. Inferring regulatory change from gene expression: the confounding effects of tissue scaling. Mol Ecol 2016; 25(20): p. 5114–5128.

82. McKenna A, Hanna M, Banks E, Sivachenko A, Cibulskis K, Kernytsky A, et al. The Genome Analysis Toolkit: a MapReduce framework for analyzing next-generation DNA sequencing data. Genome Res 2010; 20(9): p. 1297–303.

83. Li H, Handsaker B, Wysoker A, Fennell T, Ruan J, Homer N, et al. The Sequence Alignment/Map format and SAMtools. Bioinformatics 2009; 25(16): p. 2078–9.

84. Nguyen LT, Schmidt HA, von Haeseler A and Minh BQ. IQ-TREE: a fast and effective stochastic algorithm for estimating maximum-likelihood phylogenies. Mol Biol Evol 2015; 32(1): p. 268–74.

85. Patterson N, Moorjani P, Luo Y, Mallick S, Rohland N, Zhan Y, et al. Ancient admixture in human history. Genetics 2012; 192(3): p. 1065–93.

86. Danecek P, Auton A, Abecasis G, Albers CA, Banks E, DePristo MA, et al. The variant call format and VCFtools. Bioinformatics 2011; 27(15): p. 2156–8.

87. Kreitman M. Methods to detect selection in populations with applications to the human. Annu. Rev. Genomics Hum. Genet. 2000; 01: p. 539–59.

88. Conesa A, Gotz S, Garcia-Gomez JM, Terol J, Talon M and Robles M. Blast2GO: a universal tool for annotation, visualization and analysis in functional genomics research. Bioinformatics 2005; 21(18): p. 3674–6.

