## Supplementary file for "Genomic variation in the tea leafhopper reveals the basis of adaptive evolution"

^4^ Department of Biological Sciences, Brock University, 1812 Sir Isaac Brock Way, St. Catharines, ON L2S 3A1, Canada (ORCID #0000-0001-7289-2675)

**1. Genome sequencing and assembly**

**1.1 Insect samples**

Insects were collected in the South Mountain located in Fujian Agriculture and Forest University and maintained in the lab by inbreeding for 12 generations. The adults were collected for further analyzed.

**1.1 Illumina short reads sequencing**

DNA was extracted using TIANamp Genomics DNA Kit (TIANGEN, Beijing, China) and sequenced on Illumina X10 platform with 150-bp reads length and insert size of 300-500 (Supplemental Table 1).

**1.2 Genome Size and heterozygosity estimation**

Genome DNA was prepared from the adults which were maintained by sibling mating for 12 generations. Illumina PE library with insert size of 300-500 was constructed and sequenced based on Illumina X-10 platform. Totally, 37 Gb Illumina X-10 were generated. The K-mer spectra, genome heterozygosity and repeat content were generated using JELLYFISH (version 1.1.10) with default parameters [1].

**1.3 Nanopore library construction and sequencing**

Genome DNA was extracted from the adult insects using TIANamp Genomics DNA Kit. For each Nanopore library, ~8 ug gDNA was size-selected (10-50 kb) with a Blue Pippin (Sage Science, Beverly, MA) and processed using the Ligation sequencing 1 D Kit (SQK-LSK108, ONT, UK) according to the instructions. Totally 19 libraries were constructed and sequenced on 19 different R9.4 FlowCells using Nanopore GridION X5 sequencer for 48 hr each at the Genome Center of Nextomics (Wuhan, China). We use ONT Albacore software (version 0.8.4) (https://github.com/Albacore/albacore) to finish the base calling (Supplemental Table 2).

**1.4 Hi-C library construction and sequencing**

We use Hi-C to assist chromosomal-scale genome assembly. The sample crosslinking was prepared as follows: ~1000 2-3 instar nymphs were cut with scissors, then 1.25 ml 37% formaldehyde was added to obtain 2% final concentration and incubated for 10 min on plate at room temperature for further crosslink. After that, the crosslink was quenched by adding 2.5 ml 2.5M glycine (incubated for 5 min at room temperature and then incubated on ice for 15 min). Finally, the samples were centrifuged at 4 ℃ with 2000 g for 10 min and the supernatant was removed. After the crosslinking, the samples were used for Hi-C library preparation and further sequenced using Illumina X10 platform (Supplemental Table 2).

**1.5 Hi-C scaffolding and chromosome assembly**

The Hi-C reads were uniquely mapped to the contig assembly we retained the reads within 500 bp *HindIII* restriction regions for next analysis. We use 3D-DNA pipeline to correct the mis-joined contigs [2]. ALLHiC pipeline was used to link the Hi-C corrected contigs into 10 pseudo-chromosomes [3]. The accuracy of the Hi-C based assembly was evaluated by chromatin contact matrix (Supplemental Figure 2).

**1.6 Mitochondrial genome assembly**

To assemble the mitochondrial sequences, we performed a reference-guided MT genome assembly strategy according to the recent published methods [4, 5]. We first selected the Nanopore reads with length ranging from 5 kb to 16 kb, and then aligned these reads to the NCBI insect mitochondrial references using minimap2 [6] and BLAST [7]. The resulting MT reads were assembled using Canu v2.0 [8], and contigs were aligned to the published reference (*Empoasca vitis* mitochondrion, NCBI accession number: NC_024838.1) [9] by BLAST, MT contig was remain with highest identity and coverage. To polished this contig, we using corrected long reads by Racon and Medaka and then FreeBayes [10] combined with bcftools [11] using short reads. The overlapping sequences at the both ends of contig were trimmed according self-alignment using nucmer [12], and the remaining sequences is the finally circularized mitochondrial genome. Moreover, annotation was performed on the GoSeq online platform using *Empoasca vitis* mitochondrion as reference and with default parameters [13].

1.7 **Repeat annotation**

A customized de novo repeat library was built using RepeatModeler (<http://www.repeatmasker.org/RepeatModeler/>), which recruits 2 repeat finding programs including RECON (version 1.08) and RepeatSout (version 1.0.5) [14]. The generated TE consensus TE sequences were imported into the RepeatMasker (version 4.05) (http://www.repeatmasker.org) to further identify the repetitive elements. The unknown repeats were classified by TEclass (version 2.1.3) [15]. Tandem Repeat Finder (TRF) (version 4.07) [16] was used to identify the tandem repeats in the genome with the parameters: 1 1 2 80 5 200 2000 –d –h.

**1.7 Sample preparation for RNA-seq**

We collected the samples of different developmental stages (eggs, nymphs and adults). Besides, we also collected the insect samples from resistance and susceptible tea plants. Toal RNA was extracted from all these samples using Trizol following the instructions and DNA contamination was removed using RNAase-Free DNase I (Takara). These extracted RNA were sequenced on Illumina X10 platform, yielding ~3 Gb data for each sample.

**2. Population Genomics**

**2.1** **SNP Calling**

A total of 57 **s**amples were collected from 4 Tea regions (Jiangnan / South Central, Jiangbei / North Central, South China and Southwest China) of China in this study. The 150-bp pair-end Illumina reads were yielded with the average coverage ranging from 20.1× to 30.6× (Table 12S). Quality filtering was performed to remove adapters and low quality bases (Q < 30). Filtered reads were aligned against the reference genome using bwa with the default parameters. Variants were detected using the GATK toolkit (version: 3.5-0-g36282e4) [17] following the best practices workflow for variant discovery. Then the IndelRealigner was used to locally realign the resulting BAM file to remove the mismatches located around small-scale deletions and insertions. Variants were called and merged using HaplotypeCaller and GenotypeGVCFs separately. This two-step approach includes quality recalibration and re-genotyping in the merged vcf file, which can ensure the variant accuracy. Meanwhile, samtools/bcftools were applied for SNPs calling using the same data with default parameters. SNPs were further filtered based on the workflow below: (1) SNPs were filtered if existed in only one of the two pipelines (GATK or SAMtools/BCFtools); (2) SNPs located in repeat regions; (3) SNPs with high (> 1000) read depth or low reads depth (< 5); (4) SNPs with missing rate > 40%; (5) non-biallelic SNPs were removed; (6) SNPs with < 5bp distance with nearby variant sites.

**Supplementary Figures**

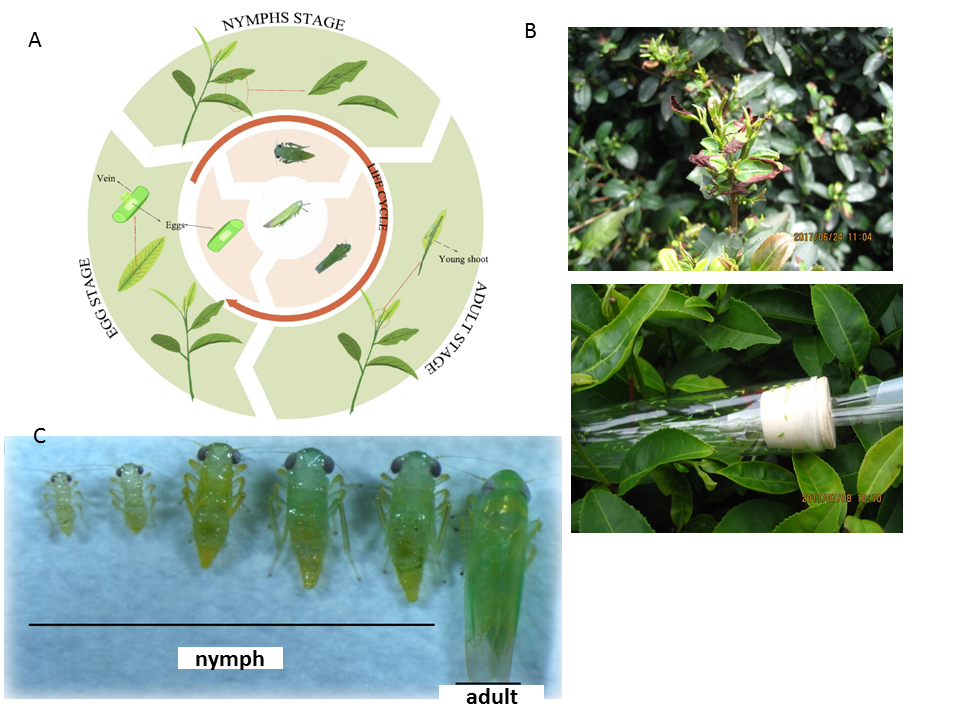

**Figure S1. Damages of *E.onukii* in modern tea plantation in China.** (A) Life cycle of *E. onukii*. (B) Damages caused by *E.onukii*. (C) Developmental stages of *E. onukii.*

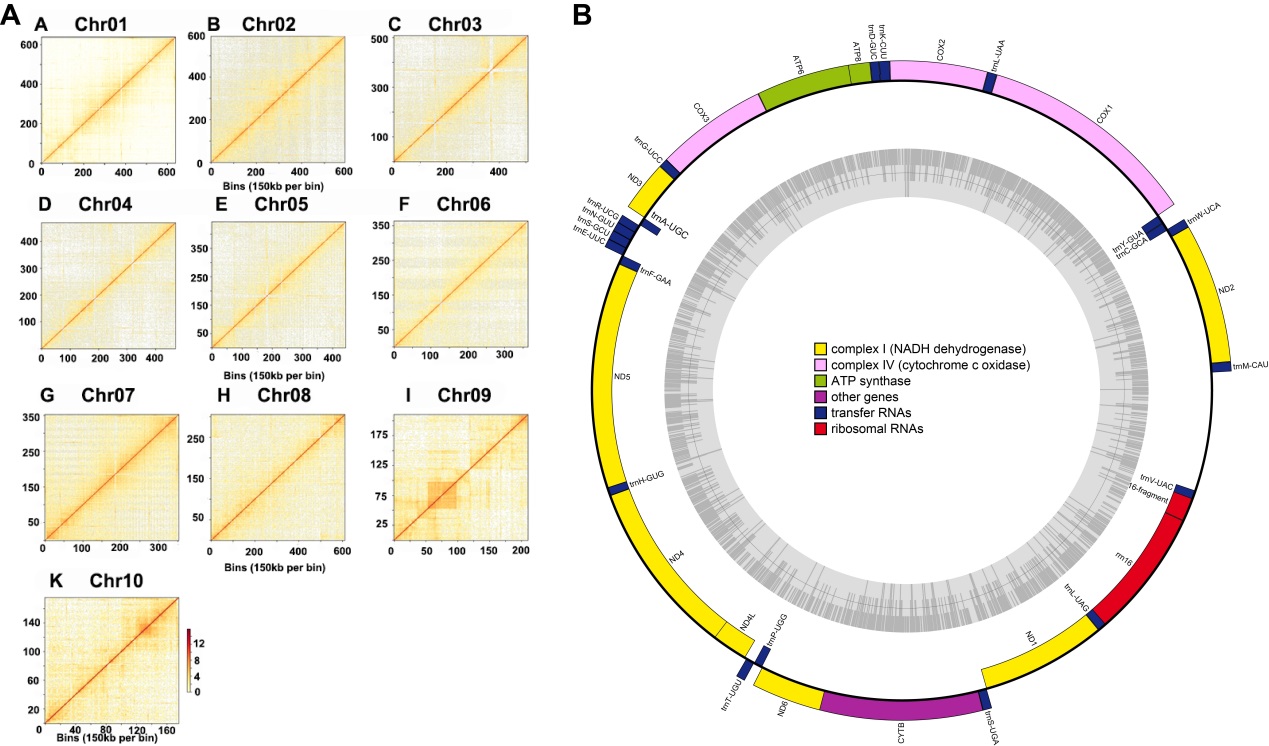

**Figure S2. Genome assembly of *E. onukii*. (A) Chromatin interactions with 150 kb resolution in *E. onukii.* (B) Mitochondrial genome of *E. onukii.*** Inner circle represents the GC content while outer circle represents genes located on mitochondrion.

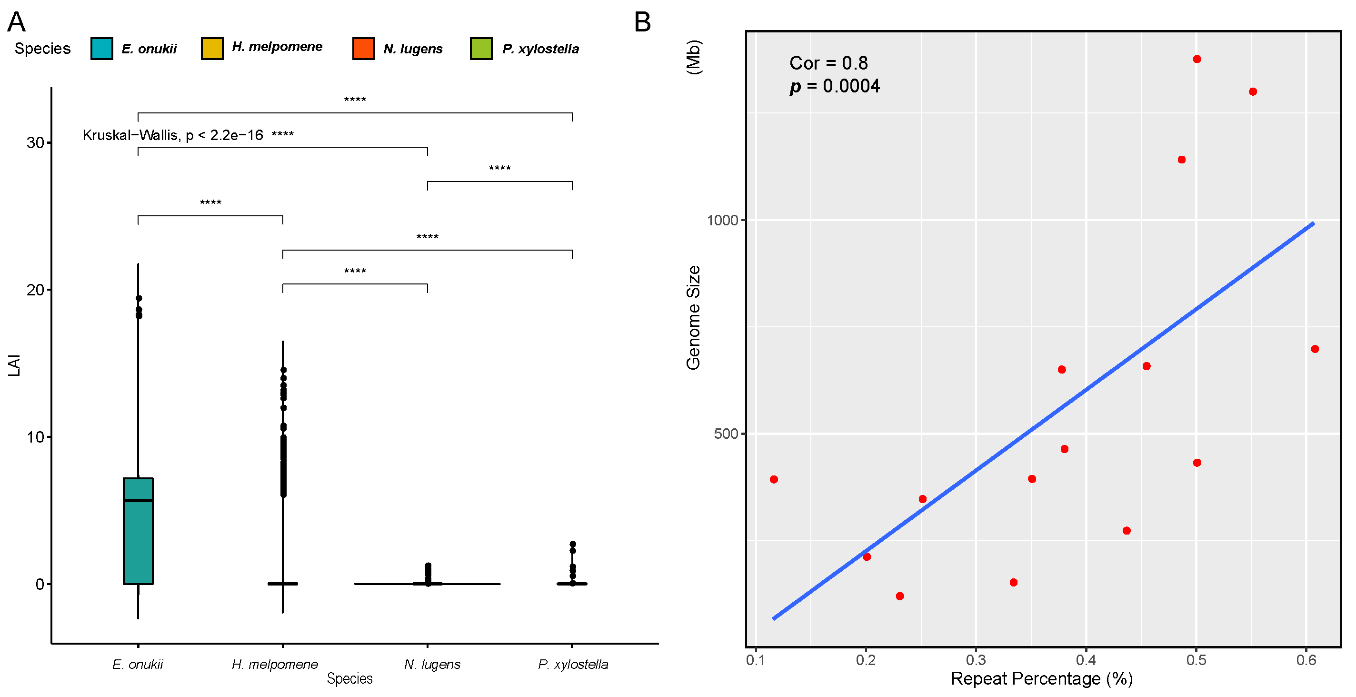

**Figure S3. (A) Assessment of the assembly using LTR Assembly Index (LAI). (B) Correlation analysis between repeat content and the genome size.**

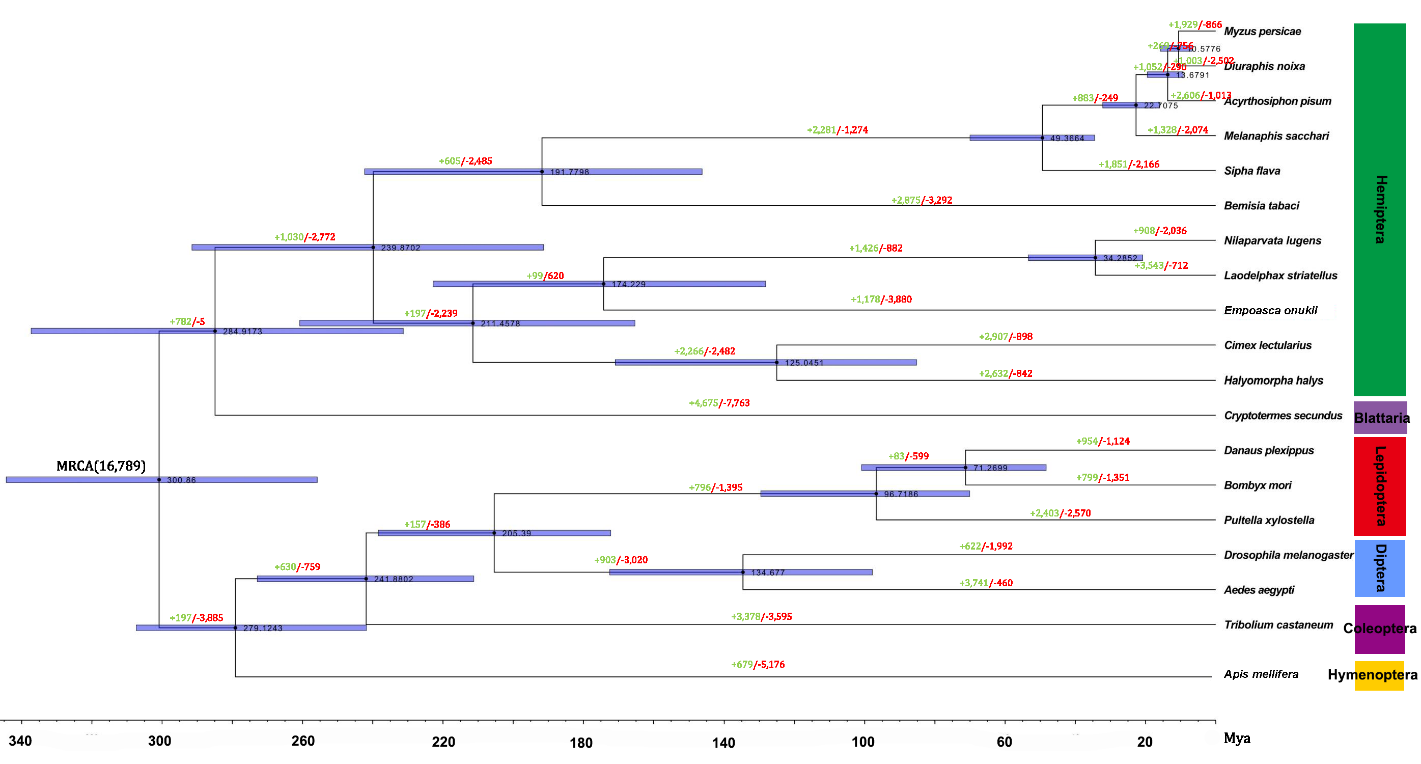

**Figure S4. Gene family expansions and contractions in the *E. onukii* compared with other insects.** Numbers for expanded (green) and contracted (red) gene families are shown on branches.

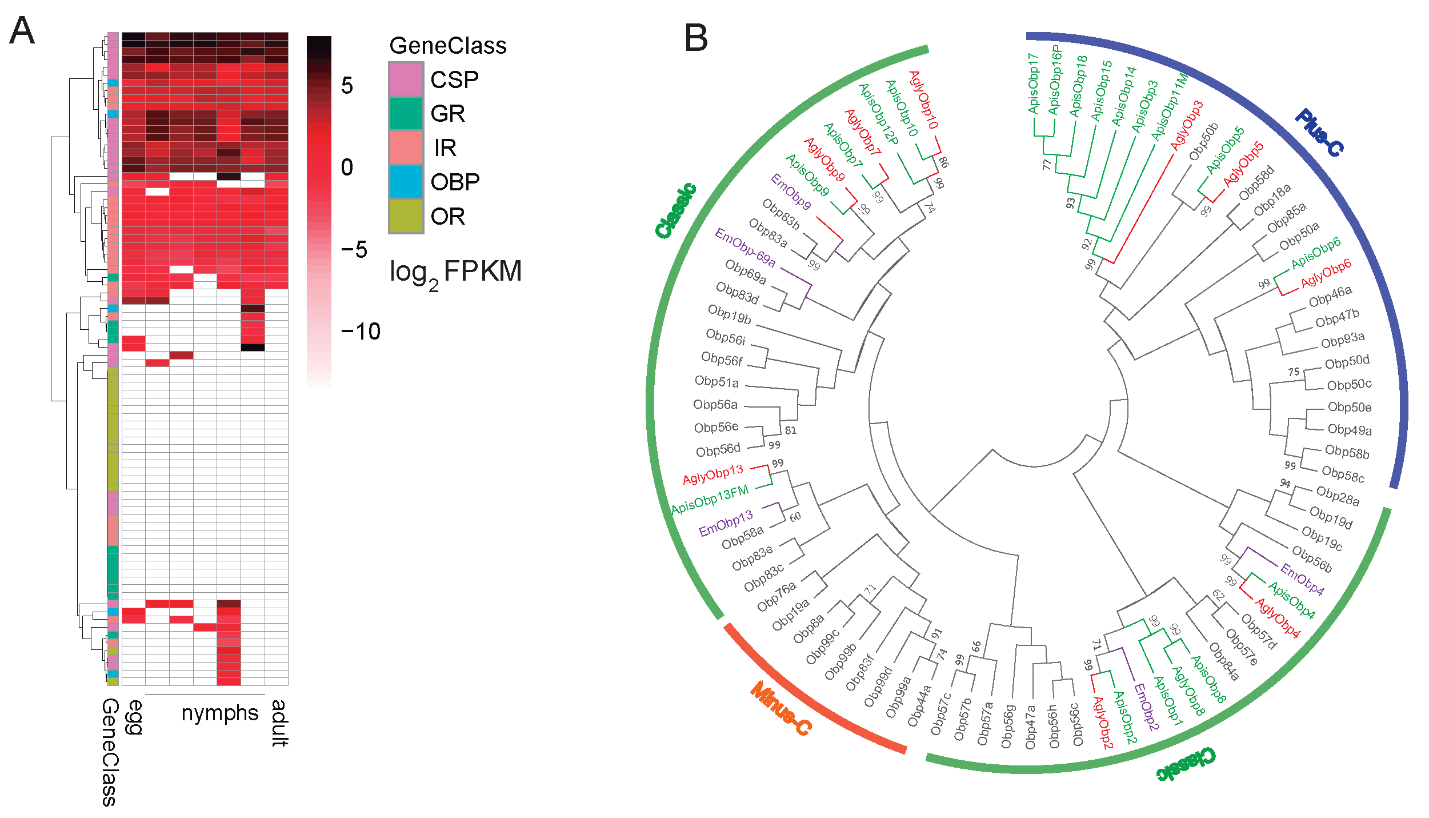

**Figure S5. Expression profile of genes involved in chemoreception and phylogenetic analysis of odorant-binding proteins (OBPs).** (A) Expression of 5 chemoreception gene families. (B) Neighbor-joining method involving 84 protein sequences was used to construct the tree. Different colors represent different species: red represents *Ap. Glycines* (Agly); green represents *A.	pisum* (Apis); purple represents *E. onukii* (Em) while black represents *D. melanogaster*.

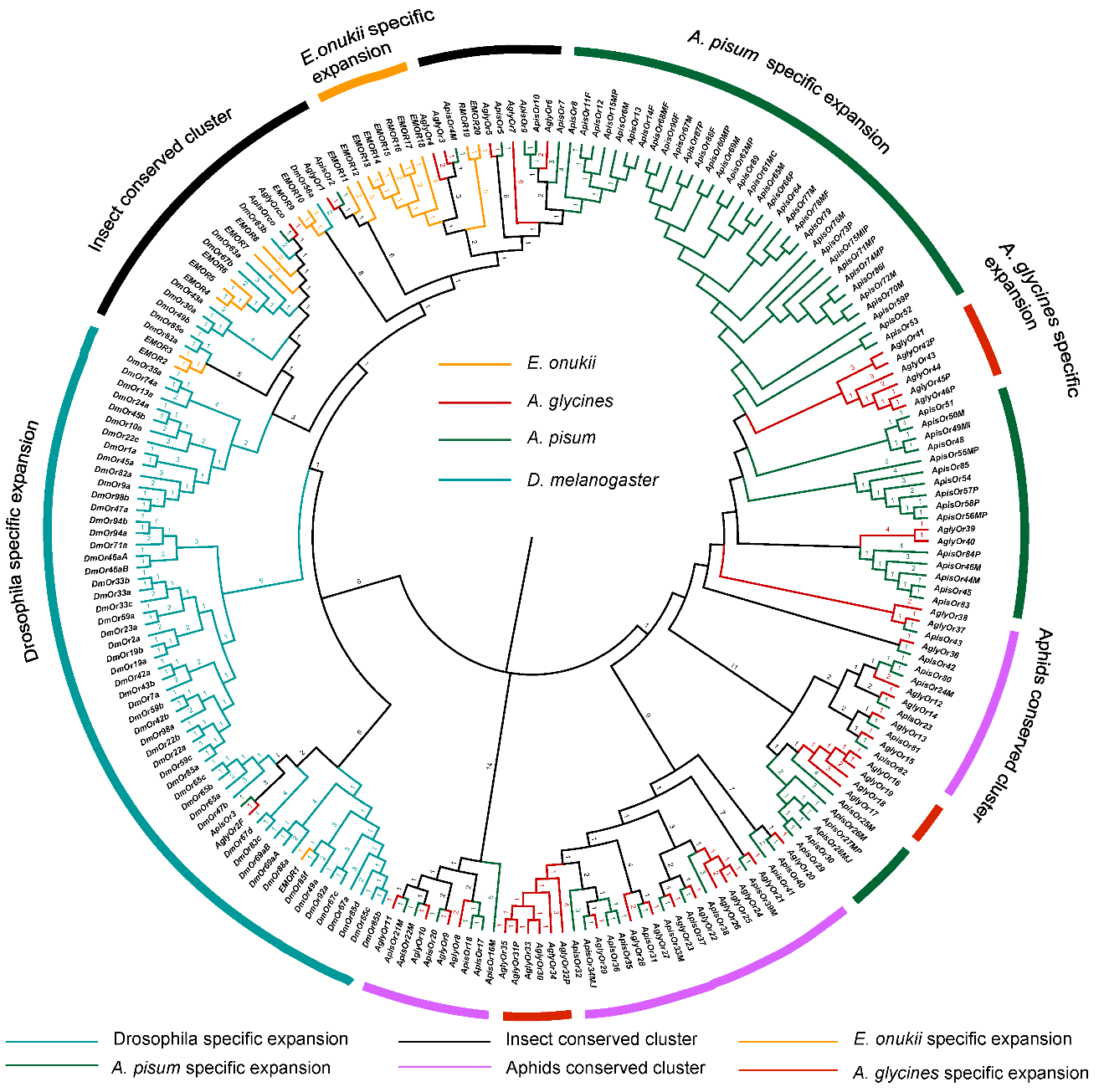

**Figure S6. Phylogenetic analysis of olfactory receptors (ORs).** Neighbor-joining method involving 216 protein sequences was used to construct the tree. Different colors represent different species: red represents *Ap. Glycines* (Agly); green represents *A.	pisum* (Apis); yellow represents *E. onukii* (Em) while blue represents *D. melanogaster*.

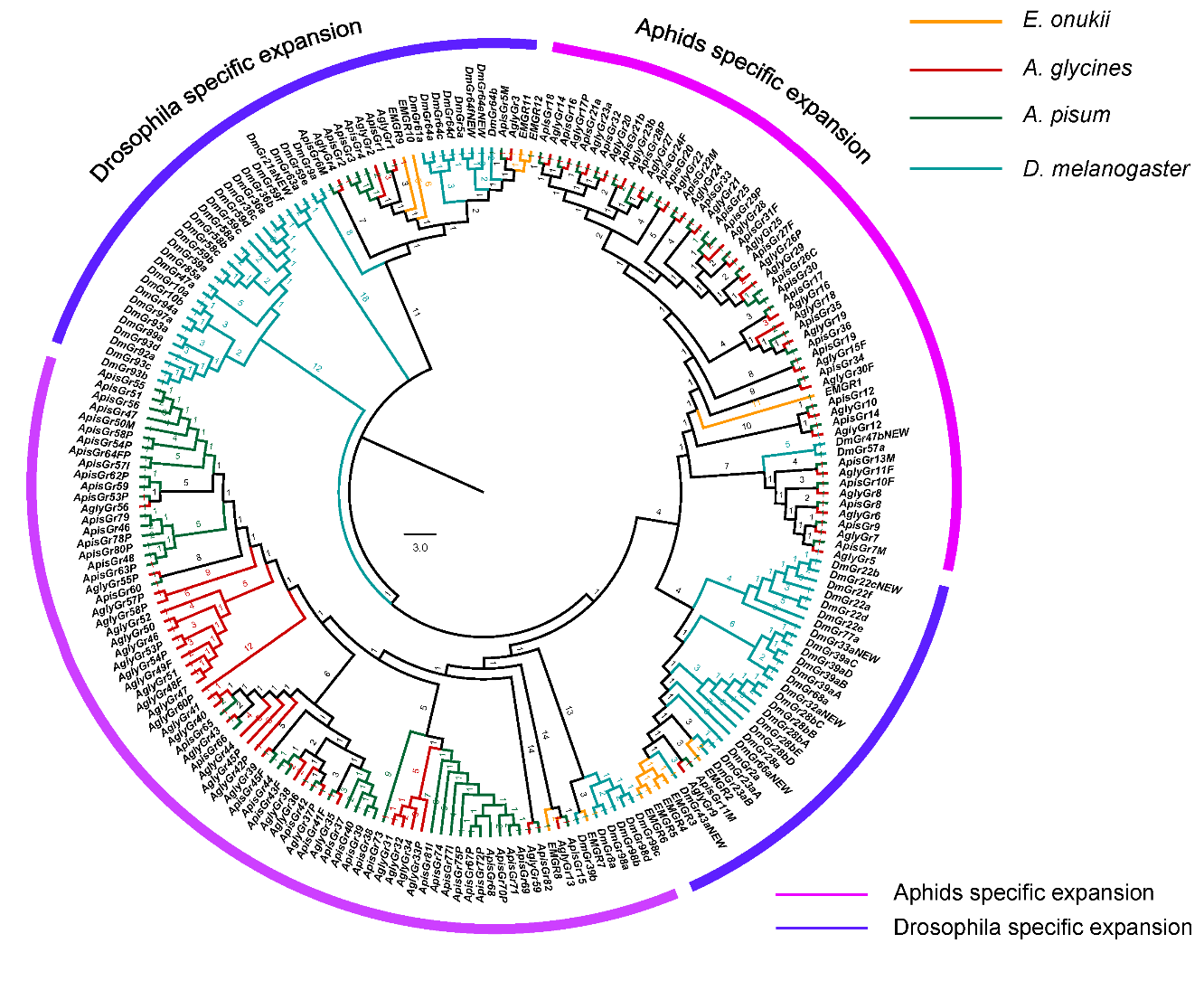

**Figure S7. Phylogenetic analysis of gustatory receptors (GRs).** Neighbor-joining method involving 219 protein sequences was used to construct the tree. Colors corresponded to Supplemental Figure 6.

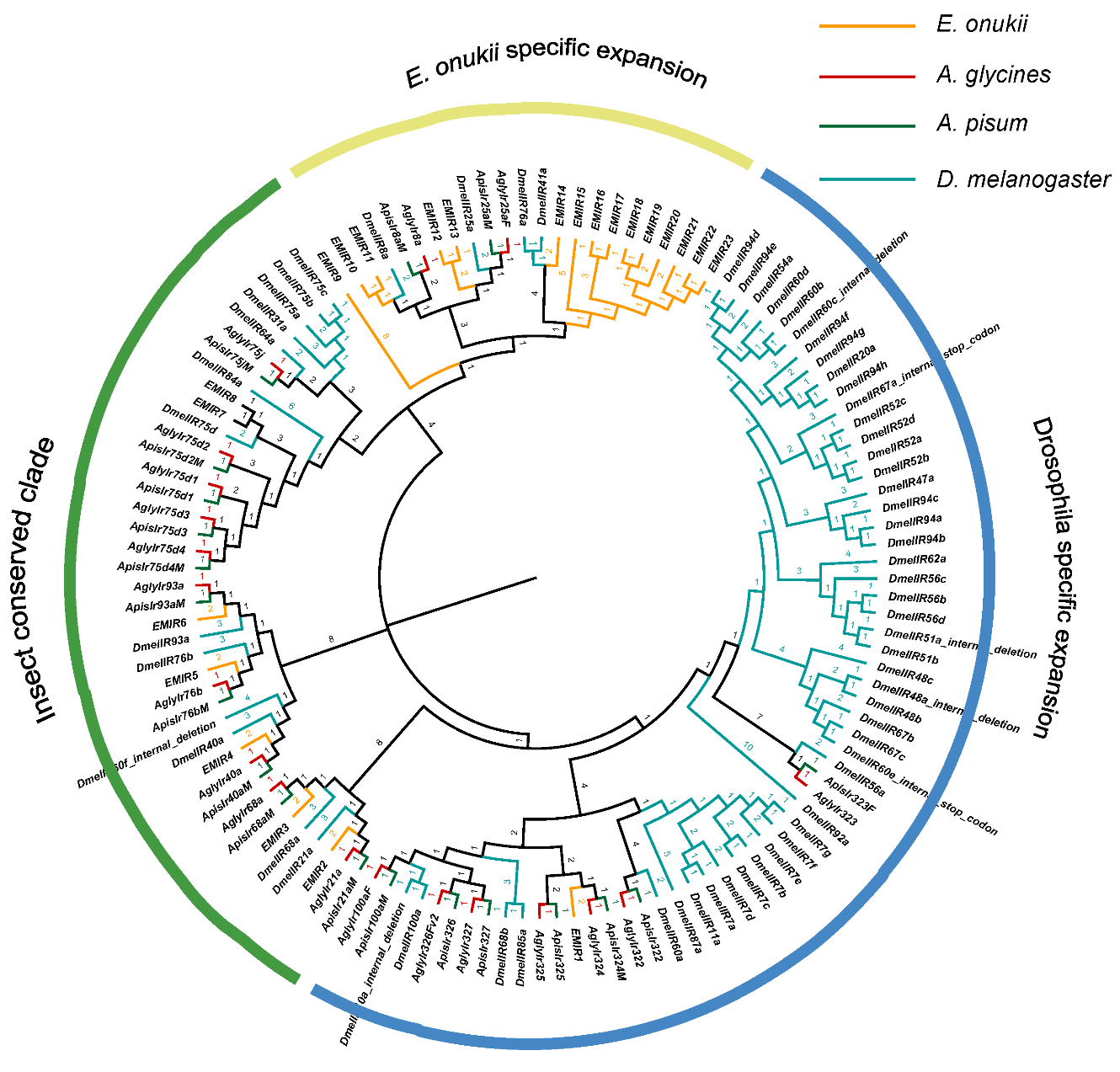

**Figure S8. Phylogenetic analysis of ionotropic receptors (IRs).** Neighbor-joining method involving 125 protein sequences was used to construct the tree. Colors corresponded to Supplemental Figure 6.

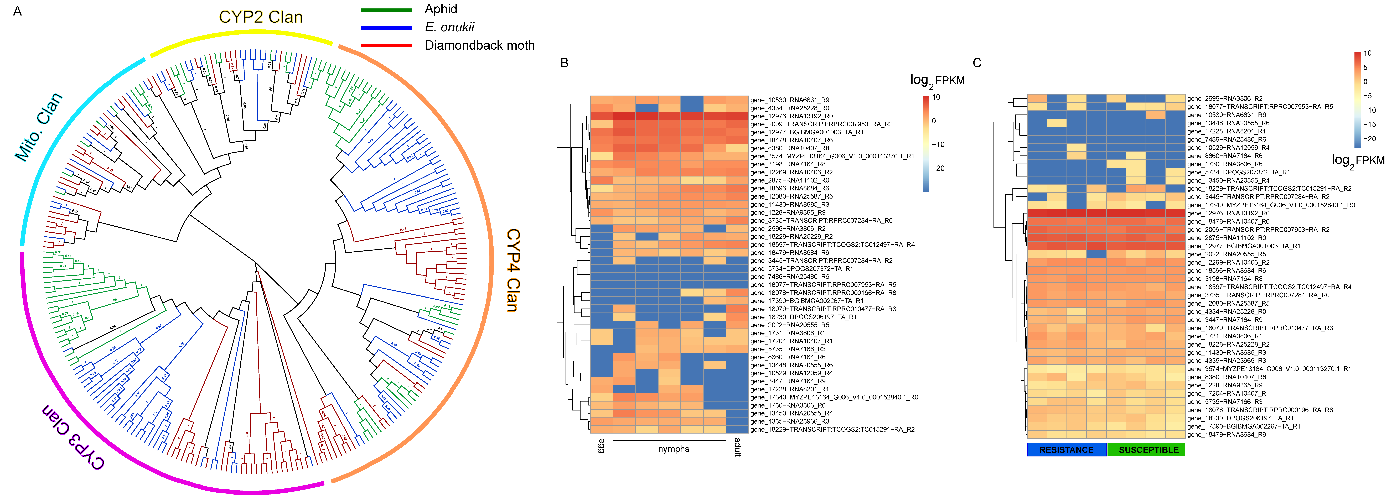

**Figure S9. Phylogenetic analysis of P450 gene family.** Neighbor-joining method involving 259 protein sequences was used to construct the tree. Colors represented different species listed in the Figure.

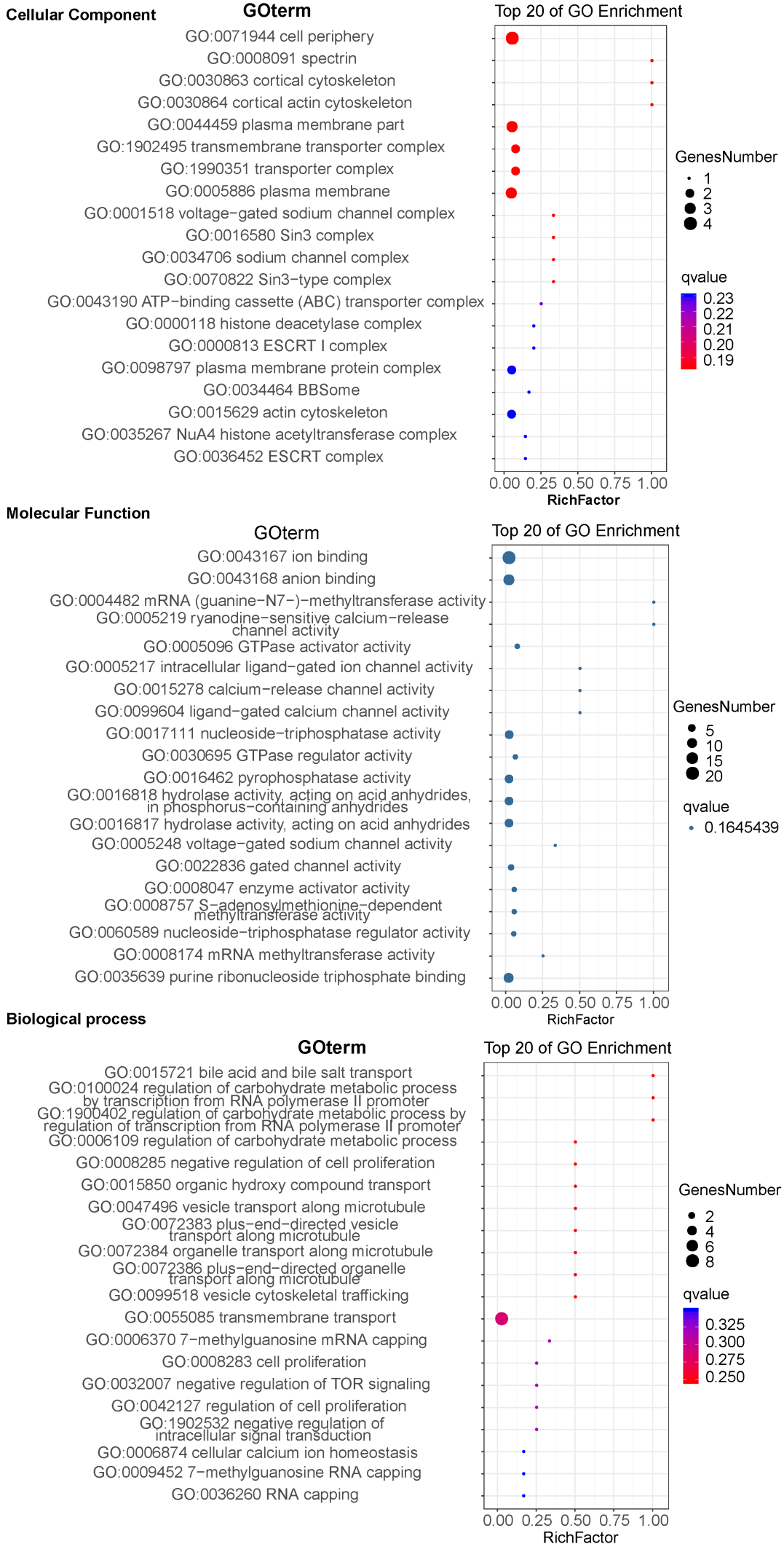

**Figure S10. GO enrichment of the genes under balancing selection in *E. onukii.* Only top 20 were listed in the figures.**

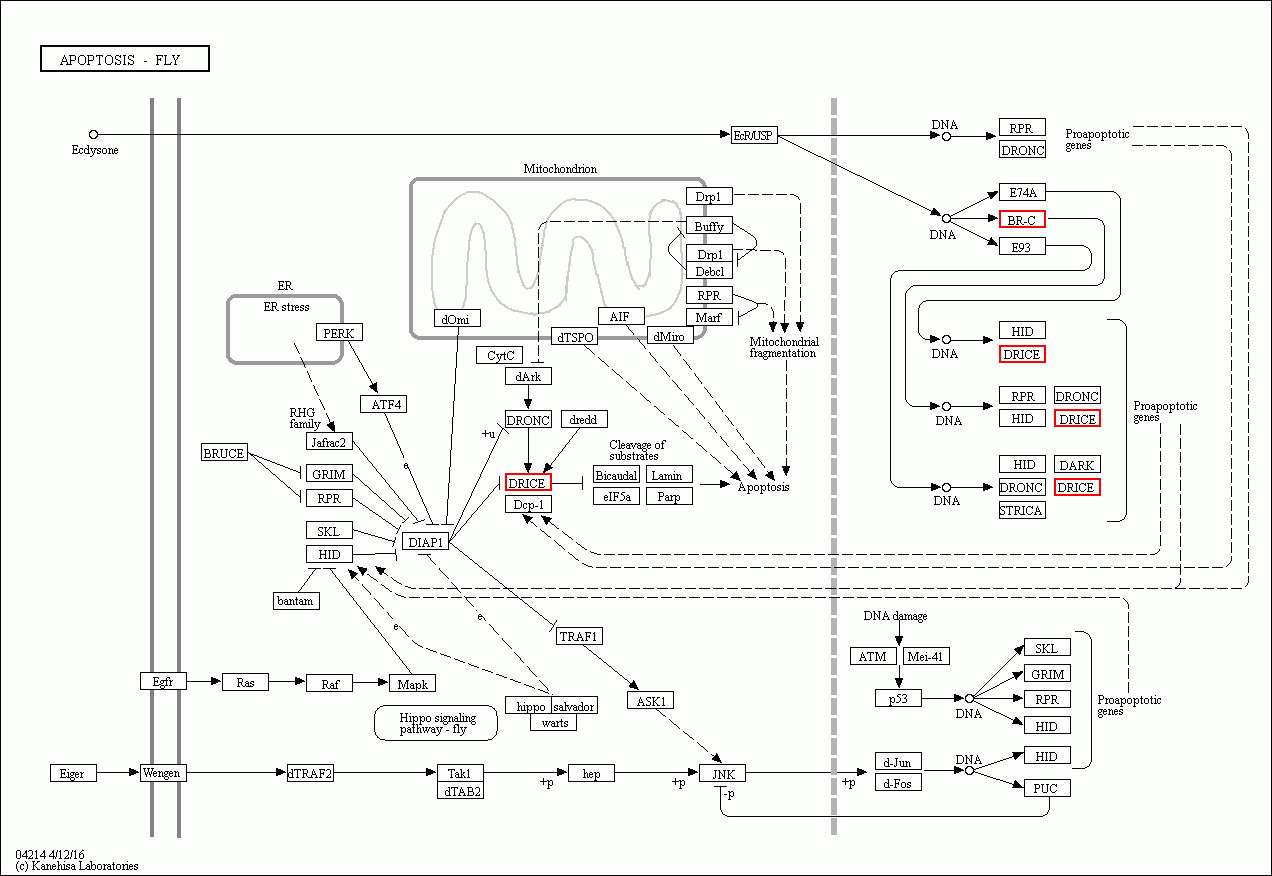

**Figure S11. Apoptosis pathway was enriched for the genes under balancing selection in *E. onukii*.** Genes under selection were marked in red.

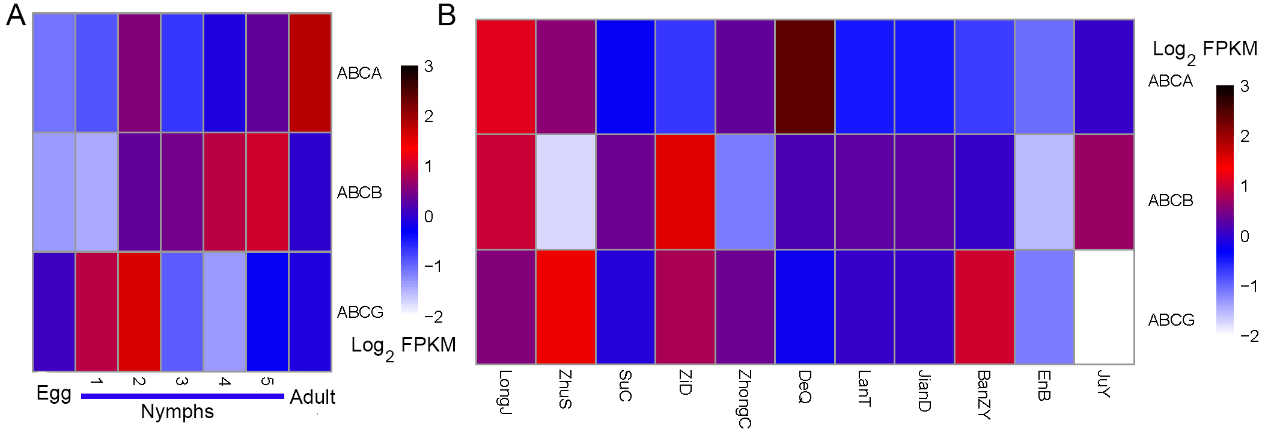

**Figure S12. Expression patterns of the ABC genes under balancing selection.** (A) Expression patterns of different developmental stages; (B) Expression patterns when living on resistant and susceptible tea cultivars.

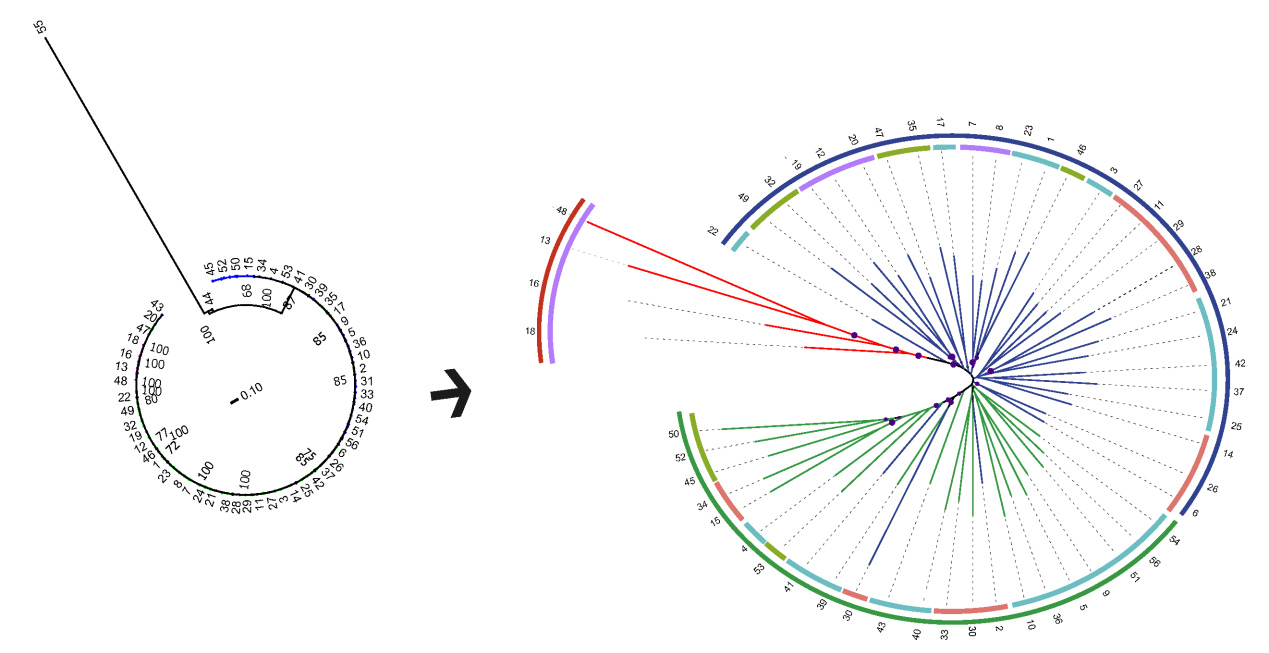

**Figure S13. Phylogenetic tree and network estimation by RAxML and SplitsTress.** Three populations according to the geographic regions that most of the individuals located: group I was Yunan (YN), Eastern China (group II); and central and southern of China (group III) were listed in Red, Green, Blue respectively (right figure) with presence of the outgroups (left figure). Different 4 tea regions of China were listed in different colors represented in Figure 3.

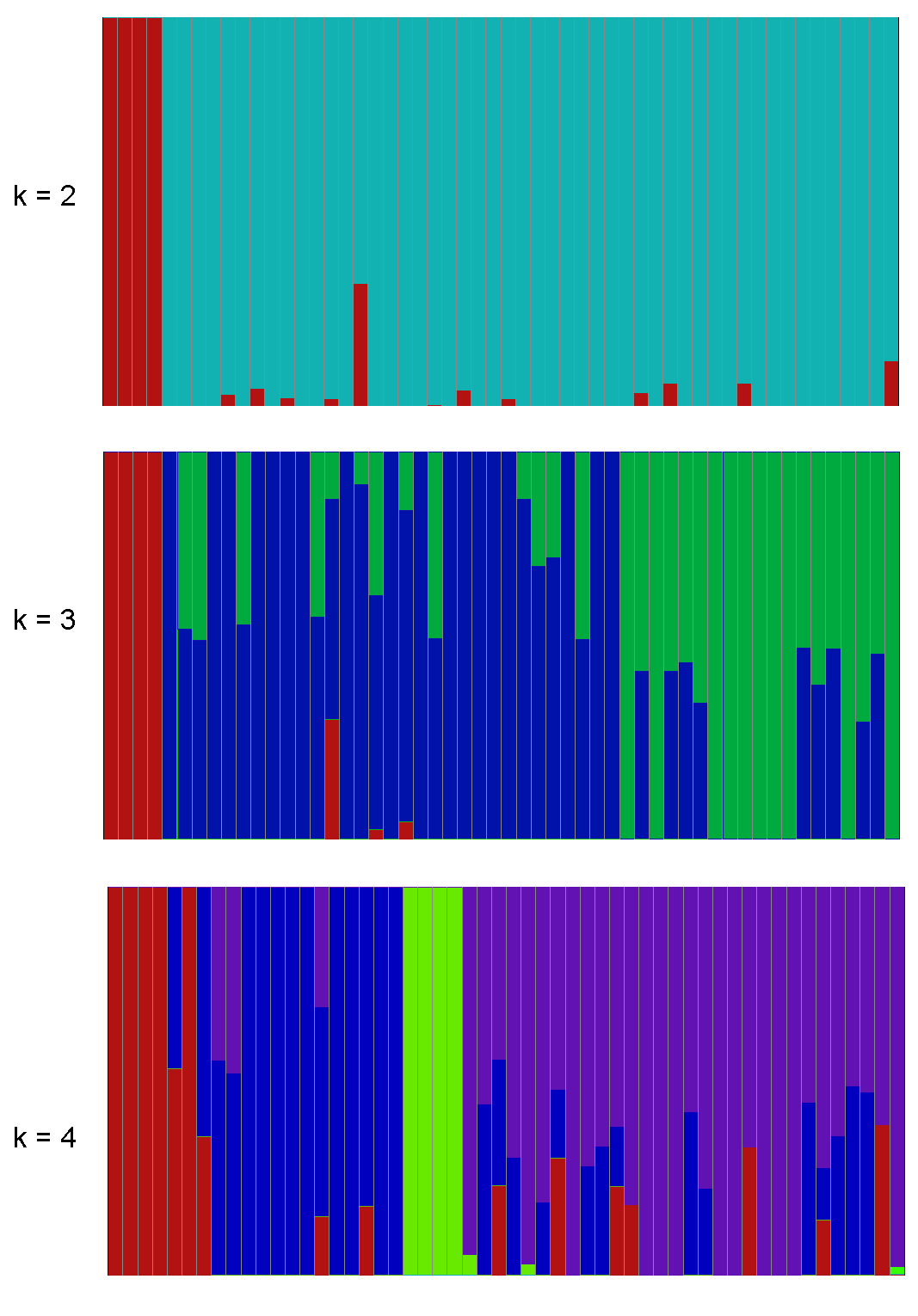

**Figure S14. Admixture analysis of 55 TGLs accessions (*k* = 2-4). IDs were represented in Supplemental Table 12.**

**Supplementary Tables**

**Table S1. Statistics of genomic sequencing data of *Empoasca onukii***

| **Items** | **Nanopore ONT** | **Illumina X10** |
| --- | --- | --- |
| Total number of reads (million) | 4 | 122 |
| Total number of sequenced bases (Gb) | 65 | 37 |
| Mean reads length (bp) | 15,737 | 150 |
| N50 (bp) | 24,965 | 150 |
| Coverage (×) | 109 | 61 |

**Table S2. Statistics of Hi-C mapping**

| **Statistics of mapping** | | |
| --- | --- | --- |
| Clean Paired-end Reads | | 123,506,761 |
| Unmapped Paired-end Reads | | 52,279,737 |
| Unmapped Paired-end Reads Rate (%) | | 42.33 |
| Paired-end Reads with Singleton | | 53,084,389 |
| Paired-end Reads with Singleton Rate (%) | | 42.98 |
| Multi Mapped Paired-end Reads | | 4,576,854 |
| Multi Mapped Ratio (%) | | 3.71 |
| Unique Mapped Paired-end Reads | | 13,565,781 |
| Unique Mapped Ratio (%) | | 10.98 |
| **Statistics of valid reads** | | |
| Unique Mapped Paired-end Reads | 13,565,781 | |
| Dangling End Paired-end Reads | 6,409,964 | |
| Dangling End Rate (%) | 42.25 | |
| Self Circle Paired-end Reads | 95,862 | |
| Self Circle Rate (%) | 0.71 | |
| Dumped Paired-end Reads | 808,192 | |
| Dumped Rate (%) | 5.96 | |
| Interaction Paired-end Reads | 5,736,909 | |
| Interaction Rate (%) | 42.29 | |
| Lib Valid Paired-end Reads | 4,225,791 | |
| Lib Valid Rate (%) | 31.15 | |
| Lib Dup (%) | 26.34 | |

**Table S3. Statistics of contig level assembly of *E. onukii***

| Items | SMART-denovo | Wtdbg2 | Quickmerge |
| --- | --- | --- | --- |
| No. of contigs | 2568 | 2420 | 1800 |
| Max length (Mb) | 2.4 | 5.7 | 12,942,971 |
| Assembly size (Mb) | 637.8 | 571.8 | 599.1 |
| N90 (bp) | 99,924 | 162,530 | 388,364 |
| N50 (bp) | 410,283 | 1,134,910 | 2,172,144 |
| Average (bp) | 248,351 | 236,266 | 332,835 |
| Complete BUSCO ratio (%) | 95.3 | 91.5 | 92.7 |
| Duplicated BUSCO ratio (%) | 3.4 | 1.4 | 2.7 |

**Table S4. BUSCO analysis of genome assembly of *E. onukii***

| **Description** | **Number** | **Percentage (%)** |
| --- | --- | --- |
| **Complete BUSCOs** | 1537 | 92.7 |
| **Complete and single-copy BUSCOs (S)** | 1493 | 90.0 |
| **Complete and duplicated BUSCOs (D)** | 44 | 2.7 |
| **Fragmented BUSCOs (F)** | 39 | 2.4 |
| **Missing BUSCOs (M)** | 82 | 4.9 |
| **Total BUSCO groups searched** | 1658 | 100 |

**Table S5. BUSCO analysis of annotation completeness**

| **Description** | **Number** | **Percentage (%)** |
| --- | --- | --- |
| **Complete BUSCOs (C)** | 1533 | 92.5 |
| **Complete and single-copy BUSCOs (S)** | 1462 | 88.2 |
| **Complete and duplicated BUSCOs (D)** | 71 | 4.3 |
| **Fragmented BUSCOs (F)** | 50 | 3.0 |
| **Missing BUSCOs (M)** | 75 | 4.5 |
| **Total BUSCO groups searched** | 1658 | 100 |

**Table S6. The statistics of different Hemiptera species assembly**

| **Species** | **Chromosome Level** | **Genome Assembly Size (Mbp)** | **Estimated Genome Size (Mbp)** | **Contig N50 (Kb)** | **Scaffold N50 (Kb)** |
| --- | --- | --- | --- | --- | --- |
| *A. pisum* | Yes | 464.3 | 512.2 | 10.8 | 22.8 |
| *A. glycines* | No | 302.9 | 317.2 | 15.8 | 174.5 |
| *D. noxia* | No | 393.0 | 417.2 | 12.6 | 397.8 |
| *M. persicae* | No | 347/355 | 409.3 | - | 435.8/164.5 |
| *N. lugens* | No | 1141 | 1220 | 24.2 | 356.6 |
| *O. fasciatus* | Yes | 1099 | 926 | 4.0 | 340.0 |
| *E. onukii* | Yes | 599.0 | 608.0 | 2200 | 67,980 |

**Table S7. Assessment of genome consistency based on NGS (Illumina) reads**

| **Items** | **Statistics** |
| --- | --- |
| Number of reads | 365,842,504 |
| Data size (Gb) | 54.88 |
| Mapped bases (Gb) | 50.64 |
| Mapping rate (%) | 92.27 |
| Genome length (Mbp) | 599 |
| Mean depth | 78.65 |
| Coverage rate (%) | 93.80 |

**Table S8. Statistics of TEs in *E. onukii* genome**

|  | **Number** | | **Length (bp)** | **% of repeats** | **% of genome** |
| --- | --- | --- | --- | --- | --- |
| **Total repeat fraction** | | 1,230,590 | 240,914,240 | 100 | 37.71 |
| **Class I: Retroelement** |  | |  |  |  |
| **LTR Retrotransposon** | 143,294 | | 25,581,399 | 10.62 | 4.00 |
| Ty1/Copia | 116 | | 90,347 | 0.04 | 0.01 |
| Ty3/Gypsy | 2801 | | 1,542,498 | 0.64 | 0.24 |
| Other | 140,377 | | 23,948,554 | 9.94 | 3.75 |
| **Non-LTR Retrotransposon** | 337,969 | | 85,834,974 | 35.63 | 13.44 |
| LINE | 242,812 | | 71,768,330 | 29.79 | 11.24 |
| SINE | 95,157 | | 14,066,644 | 5.84 | 2.20 |
| **Unclassified retroelement** | 154,284 | | 26,124,809 | 10.84 | 4.09 |
| **Class II: DNA transposon** | 383,812 | | 81,880,593 | 33.99 | 12.82 |
| **TIR** |  | |  |  |  |
| CMC [DTC] | - | | - | 0 | 0 |
| hAT | 8857 | | 1,563,092 | 0.65 | 0.24 |
| Mutator | 353 | | 80,791 | 0.03 | 0.01 |
| Tc1/Mariner | 30,632 | | 9,167,282 | 3.81 | 1.44 |
| PIF/Harbinger | 5459 | | 756,136 | 0.31 | 0.12 |
| Other | 307,879 | | 61,146,010 | 25.38 | 9.57 |
| **Helitron** | 15,028 | | 3,675,120 | 1.53 | 0.58 |
| **Tandem Repeats** | 100,545 | | 20,953,073 | 8.70 | 3.28 |
| **Unknown** | 49,267 | | 10,599,818 | 4.40 | 1.66 |

**Table S10. Gene family contraction analysis on *E.onukii* branch**

| Pfam domain | Number of genes in each species of hemipteran | | | | |
| --- | --- | --- | --- | --- | --- |
|  | *N. lugens* | *A. pisum* | *B. t* *abaci* | *E. onukii* | *M. persicae* |
| Immunoglobulin | 250 | 365 | 295 | 205 | 335 |
| Myosin | 78 | 92 | 84 | 22 | 93 |
| Tropomyosin | 22 | 31 | 50 | 10 | 30 |

**Table S13. Geographic distributions of the collected samples around China**

| **IDs** | **Location** | **Province** | **Longitude** | **Latitude** | **Tea regions** | **Clean data** | **Coverage** | **Mapping rate** | **SNPs** | **Indels** |
| --- | --- | --- | --- | --- | --- | --- | --- | --- | --- | --- |
| **1** | ShaoYang | HuNan | 110.4883333 | 27.05944444 | SYR | 13.0G | 21.8X | 98.21% | 15,154,757 | 4,280,186 |
| **2** | Wuyishan | FuJian | 118.0017889 | 27.71386389 | SER | 14.1G | 23.7X | 98.03% | 16,212,801 | 4,648,417 |
| **3** | SuiChuan | JiangXi | 114.2602583 | 26.09143611 | SYR | 13.9G | 23.3X | 98.14% | 16,130,403 | 4,622,915 |
| **4** | Sheshan | ShangHai | 121.1866306 | 31.09467222 | SYR | 14.4G | 24.2X | 98.25% | 15,858,581 | 4,530,022 |
| **5** | WuYi | ZheJiang | 119.7913472 | 28.89059444 | SYR | 16.0G | 26.9X | 98.17% | 15,891,379 | 4,589,949 |
| **6** | AnXi | FuJian | 117.8739722 | 25.00241667 | SER | 15.2G | 25.5X | 98.22% | 16,809,151 | 4,845,517 |
| **7** | DouJun | GuiZhou | 107.4756139 | 26.35382222 | SWR | 16.0G | 26.9X | 98.21% | 16,511,092 | 4,760,991 |
| **8** | FengGang | GuiZhou | 107.6997417 | 28.02914444 | SWR | 15.2G | 25.5X | 98.20% | 16,802,849 | 4,831,110 |
| **9** | LinHai | ZheJiang | 121.141831 | 28.977566 | SYR | 15.7G | 26.4X | 98.26% | 16,669,852 | 4,767,208 |
| **10** | XiHu | ZheJiang | 120.0911333 | 30.18467222 | SYR | 13.7G | 23.0X | 94.71% | 15,232,069 | 4,291,995 |
| **11** | NanNing | GuangXi | 109.1472528 | 22.75567778 | SER | 16.2G | 27.2X | 98.65% | 16,029,598 | 4,601,254 |
| **12** | PuJiang | SiChuan | 103.3807472 | 30.16399444 | SWR | 15.7G | 26.4X | 97.86% | 15,893,642 | 4,553,711 |
| **13** | TengChong | YunNan | 98.67929167 | 24.92432222 | SWR | 15.7G | 26.4X | 98.21% | 12,215,935 | 3,516,260 |
| **14** | WuZhiShan | HeNan | 109.5899972 | 18.68604167 | SER | 16.8G | 28.2X | 98.67% | 16,016,362 | 4,610,438 |
| **15** | GanZhou | GuangDong | 116.7000694 | 23.80833611 | SER | 15.8G | 26.5X | 98.32% | 14,877,178 | 4,294,313 |
| **16** | PuEr | YunNan | 100.9223944 | 22.76429167 | SWR | 18.2G | 30.6X | 90.75% | 14,011,492 | 4,010,930 |
| **17** | ShiYan | HuNan | 110.4883333 | 27.05944444 | SYR | 16.7G | 28.0X | 98.31% | 16,623,326 | 4,772,514 |
| **18** | YuXi | YunNan | 102.2699194 | 24.146275 | SWR | 15.2G | 25.5X | 98.04% | 14,805,955 | 4,235,716 |
| **19** | MoTuo | XiZang | 95.35646667 | 29.192725 | SWR | 14.7G | 24.7X | 98.38% | 15,182,502 | 4,340,031 |
| **20** | MianYang | SiChuan | 104.4594361 | 31.81407222 | SWR | 14.7G | 24.7X | 98.26% | 15,233,861 | 4,346,004 |
| **21** | ChangSha | HuNan | 113.2686111 | 28.30611111 | SYR | 13.7G | 23.0X | 97.80% | 15,334,791 | 4,365,402 |
| **22** | GuZhang | HuNan | 109.8780556 | 28.62 | SYR | 14.5G | 24.3X | 98.33% | 15,902,090 | 4,546,546 |
| **23** | EnShi | HuBei | 109.5027778 | 30.06444444 | SYR | 15.8G | 26.5X | 98.04% | 16,180,528 | 4,626,792 |
| **24** | WuHan | HuBei | 114.5027778 | 30.99611111 | SYR | 16.5G | 27.7X | 98.25% | 16,269,457 | 4,657,863 |
| **25** | LuShan | JiangXi | 115.875 | 29.50486389 | SYR | 15.7G | 26.4X | 98.37% | 16,554,166 | 4,754,175 |
| **26** | BeiFeng | FuJian | 119.3872528 | 26.153125 | SER | 16.7G | 28.0X | 98.31% | 17,021,243 | 4,898,709 |
| **27** | QingYuan | GuangDong | 113.2948111 | 24.33394722 | SER | 15.3G | 25.7X | 97.48% | 16,020,286 | 4,592,591 |
| **28** | GuiLin | GuangXi | 110.3541167 | 25.29503611 | SER | 15.7G | 26.4X | 98.28% | 15,905,714 | 4,557,483 |
| **29** | ZhanJiang | GuangDong | 110.2488778 | 20.51934444 | SER | 15.5G | 26.0X | 98.29% | 15,711,797 | 4,485,127 |
| **30** | BaiSe | GuangXi | 106.580225 | 24.46293889 | SER | 15.9G | 26.7X | 97.49% | 13,320,883 | 3,707,926 |
| **31** | FuDing | FuJian | 120.1327528 | 27.22771389 | SER | 12.2G | 20.5X | 98.43% | 15,015,367 | 4,221,552 |
| **32** | XiXiang | Shaanxi | 107.6333667 | 32.89587222 | NYR | 14.1G | 23.7X | 96.93% | 15,117,055 | 4,304,592 |
| **33** | XinZhu | TaiWan | 121.0275 | 25.19277778 | SER | 16.2G | 27.2X | 98.17% | 15,739,439 | 4,524,321 |
| **34** | NanTou | TaiWan | 121.0502778 | 24.05416667 | SER | 15.0G | 25.2X | 98.34% | 14,478,659 | 4,165,842 |
| **35** | XiShan | NeNan | 111.0791361 | 33.47525278 | NYR | 13.5G | 22.7X | 97.63% | 15,381,268 | 4,364,716 |
| **36** | NanChang | JiangXi | 116.0043611 | 28.37004722 | SYR | 15.4G | 25.9X | 98.56% | 16,502,613 | 4,703,669 |
| **37** | ZhouShan | ZheJiang | 122.0748472 | 30.08829167 | SYR | 13.3G | 22.3X | 98.35% | 15,396,559 | 4,367,697 |
| **38** | HaiKou | HaiNan | 110.4597611 | 19.46704444 | SER | 16.0G | 26.9X | 98.26% | 15,219,466 | 4,369,003 |
| **39** | LianYunGang | JiangSu | 119.3177167 | 34.66125 | SYR | 15.1G | 25.3X | 98.39% | 15,988,727 | 4,558,834 |
| **40** | QiMen | AnHui | 117.5133139 | 29.84614167 | SYR | 13.3G | 22.3X | 97.36% | 15,055,123 | 4,276,018 |
| **41** | FengYang | AnHui | 117.8724722 | 32.65033611 | SYR | 17.3G | 29.0X | 97.62% | 16,678,678 | 4,776,883 |
| **42** | LiuAn | AnHui | 116.2199889 | 31.39157778 | SYR | 14.3G | 24.0X | 98.57% | 15,956,534 | 4,535,057 |
| **43** | KaiHua | ZheJiang | 118.2636028 | 29.11968333 | SYR | 16.9G | 28.4X | 98.66% | 16,037,216 | 4,599,723 |
| **44** | Nairobi | Africa | 36.49 | 1.17 | Africa | 13.1G | 22.0X | 33.89% | 29,019 | 4749 |
| **45** | JiYuan | HeNan | 112.4295694 | 35.191525 | NYR | 13.8G | 23.2X | 98.57% | 14,738,189 | 4,185,898 |
| **46** | XinYang | HeNan | 113.7857583 | 32.19541389 | NYR | 12.4G | 20.8X | 98.56% | 15,018,422 | 4,238,150 |
| **47** | LongNan | GanSu | 105.2887 | 32.73546944 | NYR | 16.1G | 27.0X | 96.25% | 15,877,287 | 4,544,245 |
| **48** | MengHai | YunNan | 100.4997083 | 21.79693889 | SWR | 13.5G | 22.7X | 98.27% | 11,055,619 | 3,150,374 |
| **49** | ZiYang | ShanXi | 108.5241639 | 32.52690833 | NYR | 15.0G | 25.2X | 92.10% | 13,869,412 | 3,950,991 |
| **50** | TaiShan | ShanDong | 117.1619167 | 36.27371667 | NYR | 18.3G | 30.7X | 80.30% | 13,580,586 | 3,815,772 |
| **51** | SuZhou | JiangSu | 120.3835833 | 31.09733333 | SYR | 20.7G | 34.7X | 91.89% | 16,483,588 | 4,744,757 |
| **52** | RiZhao | ShanDong | 119.2616833 | 35.31455 | NYR | 15.7G | 26.4X | 92.71% | 14,785,789 | 4,197,553 |
| **53** | LaoShan | ShanDong | 120.6853333 | 36.15211667 | NYR | 14.5G | 24.3X | 98.60% | 15,455,293 | 4,399,335 |
| **54** | TaiZHou | ZheJiang | 121.141831 | 28.977566 | SYR | 16.9G | 28.4X | 98.43% | 16,836,234 | 4,835,103 |
| **55** | Toronto | Canada | 79.25 | 43.4 | Canada | 14.6G | 24.5X | 39.17% | 375,579 | 63,563 |
| **56** | LiShui | ZheJiang | 119.3290116 | 28.035737 | SYR | 15.0G | 25.2X | 98.33% | 16,439,489 | 4,685,444 |

**Table S16. Functional analysis of genes under purifying selection**

| **Gene ID** | **Functional annotation** | **E-value** |
| --- | --- | --- |
| gene_10670-MYZPE13164_G006_V1.0_000135270.2_R1 | Myelin regulatory factor | 0 |
| gene_11818-RNA12909_R0 | Forkhead box protein P1 | 9e-151 |
| gene_15492-RNA10461_R0 | Coronin-6 | 0 |
| gene_15488-RNA19235_R0 | Inositol hexakisphosphate and Diphosphoinositol-pentakisphosphate kinase 2 | 0 |
| gene_15510-RNA2388_R0 | KICSTOR complex protein SZT2-like | 0 |
| gene_15507-RNA5810_R0 | Protein tyrosine phosphatase | 0 |
| gene_15477-GB41265-RA_R0 | Nucleoredoxin-like isoform | 2e-75 |

**Reference**

[1].	Marcais G and Kingsford C. A fast, lock-free approach for efficient parallel counting of occurrences of k-mers*.* Bioinformatics 2011; 27(6): p. 764-70.

[2].	Dudchenko O, Batra SS, Omer AD, Nyquist SK, Hoeger M, Durand NC, et al. De novo assembly of the Aedes aegypti genome using Hi-C yields chromosome-length scaffolds*.* Science 2017; 356(6333): p. 92-95.

[3].	Zhang X, Zhang S, Zhao Q, Ming R and Tang H. Assembly of allele-aware, chromosomal-scale autopolyploid genomes based on Hi-C data*.* Nat Plants 2019; 5(8): p. 833-845.

[4].	Masterson P, Clark K, Martin F, Howe K, Flicek P, Walenz BP, et al. Towards complete and error-free genome assemblies of all vertebrate species*.* Nature 2021; 592(7856): p. 737-746.

[5].	Formenti G, Rhie A, Balacco J, Haase B, Mountcastle J, Fedrigo O, et al. Complete vertebrate mitogenomes reveal widespread repeats and gene duplications*.* Genome Biol 2021; 22(1): p. 120.

[6].	Li H. Minimap2: pairwise alignment for nucleotide sequences*.* Bioinformatics 2018; 34(18): p. 3094-3100.

[7].	Altschul SF, Gish W, Miller W, Myers EW and Lipman DJ. Basic local alignment search tool*.* J Mol Biol 1990; 215(3): p. 403-10.

[8].	Koren S, Walenz BP, Berlin K, Miller JR, Bergman NH and Phillippy AM. Canu: scalable and accurate long-read assembly via adaptive k-mer weighting and repeat separation*.* Genome Res 2017; 27(5): p. 722-736.

[9].	Zhou N, Wang M, Cui L, Chen X and Han B. Complete mitochondrial genome of Empoasca vitis (Hemiptera: Cicadellidae)*.* Mitochondrial DNA A DNA Mapp Seq Anal 2016; 27(2): p. 1052-3.

[10].	Garrison E and Marth G. Haplotype-based variant detection from short-read sequencing*.* arXiv:1207.3907 [q-bio] 2012.

[11].	Danecek P, Auton A, Abecasis G, Albers CA, Banks E, DePristo MA, et al. The variant call format and VCFtools*.* Bioinformatics 2011; 27(15): p. 2156-8.

[12].	Marcais G, Delcher AL, Phillippy AM, Coston R, Salzberg SL and Zimin A. MUMmer4: A fast and versatile genome alignment system*.* PLoS Comput Biol 2018; 14(1): p. e1005944.

[13].	Tillich M, Lehwark P, Pellizzer T, Ulbricht-Jones ES, Fischer A, Bock R, et al. GeSeq - versatile and accurate annotation of organelle genomes*.* Nucleic Acids Res 2017; 45(W1): p. W6-W11.

[14].	Price AL, Jones NC and Pevzner PA. De novo identification of repeat families in large genomes*.* Bioinformatics 2005; 21 Suppl 1: p. i351-8.

[15].	Abrusan G, Grundmann N, DeMester L and Makalowski W. TEclass--a tool for automated classification of unknown eukaryotic transposable elements*.* Bioinformatics 2009; 25(10): p. 1329-30.

[16].	Benson G. Tandem repeats finder: a program to analyze DNA sequences*.* Nucleic Acids Res 1999; 27(2): p. 573-80.

[17].	McKenna A, Hanna M, Banks E, Sivachenko A, Cibulskis K, Kernytsky A, et al. The Genome Analysis Toolkit: a MapReduce framework for analyzing next-generation DNA sequencing data*.* Genome Res 2010; 20(9): p. 1297-303.
